# Actomyosin Contractility is a Potent Suppressor of Mesoderm Induction by Human Pluripotent Stem Cells

**DOI:** 10.1101/2024.09.30.615859

**Authors:** Loic Fort, Vaishna Vamadevan, Wenjun Wang, Ian G. Macara

## Abstract

The activation of WNT signaling in human pluripotent stem cells (hPSCs) drives efficient conversion to lateral mesoderm, which can be further differentiated into cardiomyocytes. Stabilization of the WNT effector β-catenin promotes expression of mesoderm-specifying genes such as *TBXT* (which encodes Brachyury) and drives an epithelial-mesenchymal transition (EMT). Mechanical forces are essential for the self-organization and development of vertebrate embryos but the role of forces, especially actomyosin contractility, in mesoderm specification has remained controversial. We discovered that, unexpectedly, increasing actomyosin contractility by expression of constitutively active Rho kinase or MLC kinase, efficiently blocked any induction of mesoderm by WNT signaling, and cells failed to undergo EMT. Conversely, the suppression of contractility by inhibitors and genetic approaches significantly accelerated differentiation: Brachyury induction was enhanced, and EMT initiated 24hrs earlier than in control settings. These data were initially puzzling because we observed that WNT signaling was sufficient by itself to promote contractility of hiPSC colonies. Notably, however, we showed that contractility must be inhibited prior to WNT activation to efficiently promote mesoderm specification, suggesting that reduced tension primes the pluripotent state, not the induced state, to accelerate differentiation along this trajectory. Mechanistically, we found that loss of contractility decreased junctional β-catenin and promoted active β-catenin levels in the cytoplasm and nucleus. Increased contractility had opposite effects, highlighting actomyosin contractility at the pluripotent state as a key regulator of WNT signaling responsiveness through effects on adherens junctions.

## Introduction

Mechanical forces play major roles at many stages of animal development: Starting as early as compaction at the 8-cell stage ^1^, mechanics plays a central role in self-organization to shape the embryo ^2–5^. In vitro, the differentiation of pluripotent stem cells (PSCs) into germ layers is governed by dynamic interplay between mechanical forces, signaling cues, and transcriptional regulators. Internal forces are largely generated by actomyosin contractility, and in epithelia are transmitted across cell boundaries by intercellular junctions, enabling collective organization ^6–11^. But how actomyosin contractility is coupled to cell fate specification through target gene expression has remained unclear.

Human embryonic stem cell (hESC) and induced pluripotent stem cell (hiPSC) colonies can be induced to select the mesoderm lineage and eventually to differentiate into functional cardiomyocytes ^12–14^ Both types of PSCs are strongly epithelial, and lineage specification relies on an epithelial-to-mesenchymal transition (EMT) that is reminiscent of some aspects of gastrulation ^15–17^.

Mesoderm can also be generated from hPSCs by direct WNT pathway activation using the small molecule GSK3β inhibitor Chiron 99021 (hereafter CHIR), which protects cytosolic β-catenin from degradation through a similar mechanism to WNT ligand signaling. The stabilized β-catenin then enters the nucleus and binds to TCF transcription factors at target gene promoters. A separate pool of β-catenin in epithelial cells is tightly associated with E-cadherin at the adherens junctions, but whether this pool contributes to WNT signaling and mesoderm induction by hPSCs, or participates in WNT responses in other situations, remains controversial ^18^. For example, increased junctional β-catenin has been reported to correlate with increased nuclear β-catenin, though the mechanism for this phenomenon was not explained ^17^. To the contrary, in *Drosophila* imaginal disks, actomyosin contractility can promote E-cadherin accumulation at adherens junctions^11^, which reduces β-catenin-mediated gene expression ^11^, while deletion of E-cadherin promotes WNT-responsive gene expression ^19^. These studies in *Drosophila* suggest that cytoplasmic β-catenin and β-catenin at the adherens junctions are dynamically linked, but whether a similar titration system applies in human cells and, particularly in hPSCs, remains unknown. Although there are multiple studies on the impact of extracellular matrix stiffness ^16,17^, tissue confinement ^20^, and EMT ^21^ on the E-cadherin/β-catenin interaction and its downstream consequences, there has been to our knowledge no direct interrogation of how actomyosin contractility impacts mesoderm commitment by hPSCs.

From prior work on BMP4-driven mesoderm in hESCs ^16,17^, we expected that inhibiting actomyosin activity would block induction. Surprisingly, however, we obtained the opposite result: increasing contractility was sufficient to completely block differentiation and EMT, while suppressing MLC phosphorylation using small molecule inhibitors or genetic tools substantially promoted mesoderm induction. These data were also puzzling because induction of differentiation alone promotes contractility of stem cell colonies. However, we discovered that the effects of elevated or suppressed actomyosin contractility have a temporal dependence, since their effects on differentiation depend on priming of the pluripotent state, not on events post-induction. Mechanistically, we found that promoting cell relaxation led to loss of β-catenin from cell junctions and an increase in active β-catenin in the cytoplasm and nucleus, suggesting that the β-catenin pools are dynamically coupled through actomyosin contractility to control mesoderm specification.

## Results

### Contractility is a potent inhibitor of mesoderm identity

We initially hypothesized that actomyosin contractility might support mesoderm specification, given prior reports implicating mechanical forces in lineage commitment ^5,16,17^. The broad-spectrum phosphatase inhibitor Calyculin A has been used in several studies of the impact of enhanced contractility on embryonic development ^5,9^ but it blocks dephosphorylation of scores of proteins, not just ppMLC. Therefore, we treated WT hiPSCs before and during differentiation with RhoA activator II (CN03), a highly specific bacterial toxin derivative that converts the RhoA Glutamine 63 to Glutamate, locking RhoA in a constitutively active form ^22,23^, which will then specifically activate ROCK to phosphorylate MLC (**Supplementary Figure 1A**). Staining for phospho-T18/S19 Myosin Light Chain 2 (ppMLC2) confirmed that CN03 treatment led to higher levels of contractility by 24 hrs post-CHIR addition (**Supplementary Figure 1B**). Strikingly, CN03-treated cells failed to express any detectable Brachyury (encoded by *TBXT* gene, also known as *T* gene), a mesoderm marker. (**Supplementary Figure 1C-D**). However, CN03 treatment caused cell toxicity past 48h, preventing further analysis.

For this reason, we turned to genetic approaches by stably expressing constitutively active ROCK2 or MLCK (ROCK2^CA^ and MLCK^CA^) under a Doxycycline inducible promoter, with constitutive expression of a Venus reporter. We first confirmed that Venus^Pos^ cells treated with Doxycycline showed signs of higher contractility. In co-cultures containing ROCK2^CA^/WT or MLCK^CA^/WT cells treated with Doxycycline, Venus^Pos^ clusters showed strong cortical F-actin and higher ppMLC2 staining organized as fibers, compared to WT Venus^Neg^ cells, which show more diffuse staining (**Supplementary Figure 1E, G**). These data support the idea that Venus^Pos^ cells experience higher contractility, allowing us to test the impact on cell fate. As an in situ control unlabeled WT hiPSCs were mixed 1:1 with green (Venus^Pos^) transduced cells, and the cultures were pre-treated with Doxycycline (or vehicle) to turn on ROCK2^CA^ or MLCK^CA^ before starting the differentiation protocol. We prolonged the differentiation to 72 hrs in this co-culture system to better assess the impact of enhanced contractility (**Figure 1A**). Venus-positive and negative areas were outlined based on Venus expression across clusters (**Supplementary Figure 1F, H**) and percentages of cells expressing mesoderm and EMT markers were measured. Strikingly, following 72h of differentiation in the presence of doxycycline, Venus^Pos^ clusters of ROCK2^CA^ cells showed a dramatic decrease in mesoderm (**Figure 1B, C**), primitive streak (**Figure 1D, E**) compared to Venus^Neg^ cells or Vehicle-treated cells. Venus^Pos^ cells also failed to undergo EMT, which usually occurs 50-52h post differentiation, as observed by the absence of Slug expression and the persistence of epithelial marker ZO-1 (**Figure 1F-H**).

**Figure 1:**
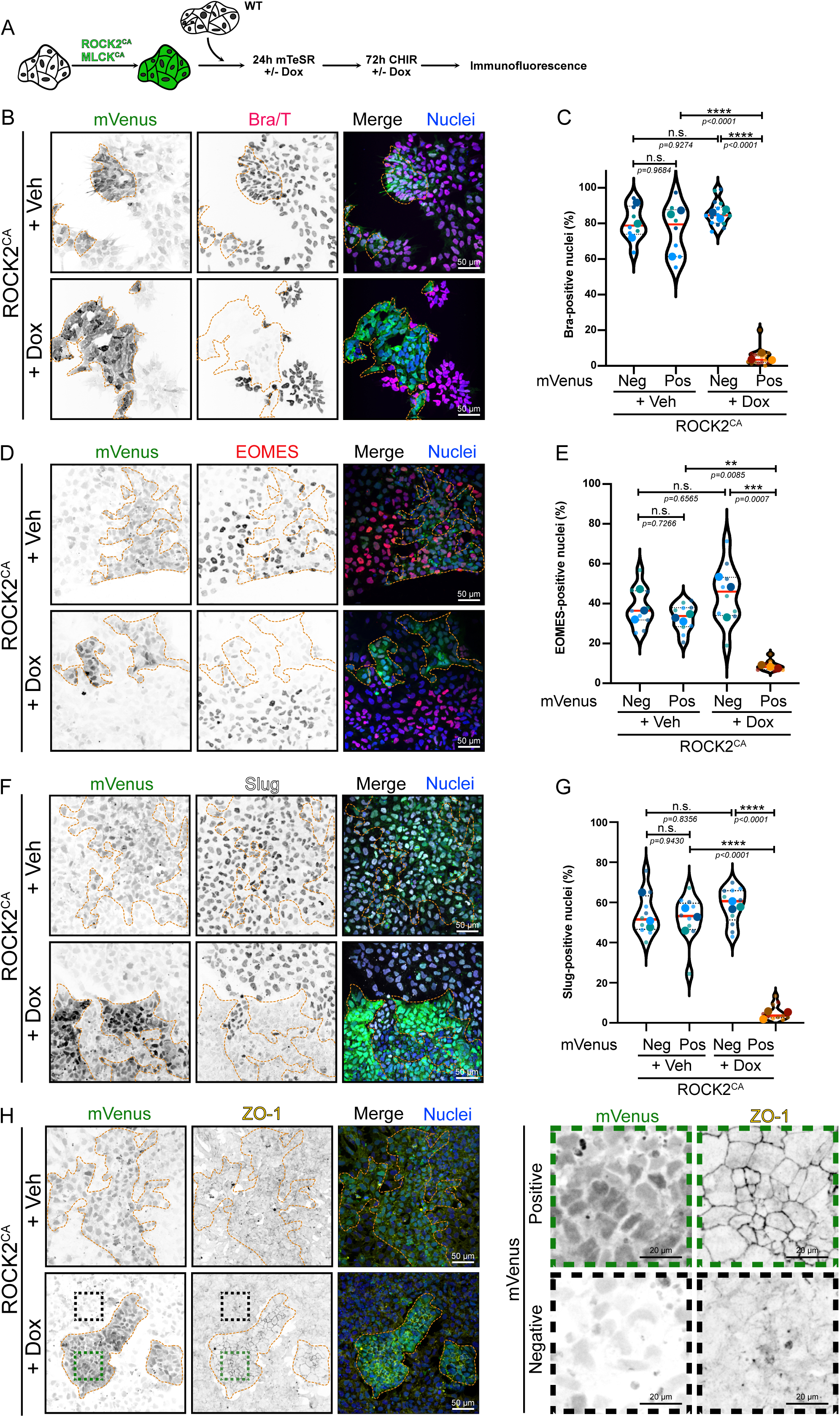
Increased actomyosin contractility is sufficient to block mesoderm commitment and EMT. (A) Experimental design for co-culture experiment. hiPSC were transduced with lentivector expressing ROCK2^CA^ or MLCK^CA^. mVenus positive-transduced cells were mixed with mVenus-negative WT hiPSCs. ROCK2^CA^/MLCK ^CA^ expression is induced by addition of Doxycycline 24h prior to initiating mesoderm differentiation. Doxycycline induction was maintained during the 3 days of differentiation using CHIR before fixing and staining. (B-H) Representative MaxIP immunofluorescence for Brachyury (B), EOMES (D), Slug (F) and ZO-1 (H) in Vehicle and Doxycycline-induced ROCK2^CA^ co-culture. mVenus-positive cell clusters are highlighted by an orange dotted line. Individual channels are presented as inverted LUT. Scale bar = 50 µm. For (H), magnified mVenus-positive and mVenus-negative areas are shown as insets. Scale bar = 20 µm. Quantification of Brachyury (C), EOMES (E) and Slug-positive cells (G) is reported as violin plot, comparing mVenus-positive *vs* mVenus-negative clusters in the presence or absence of Doxycycline. Median (Plain red line) and quartiles (Dotted black lines). For Brachyury, n = 7 (+ Veh) and n=15 (+ Dox) across N=3 independent biological repeats. For EOMES, n=9 across N=3 independent biological repeats. For Slug, n=9 (+ Veh) and n=10 (+ Dox) across N=3 independent biological repeats. One-way ANOVA with Tukey’s multiple comparisons post-test was performed.

In addition to ROCK2, MLC2 phosphorylation was previously shown to be spatially regulated by Myosin Light Chain Kinase (MLCK) in polarized and non-polarized cells ^24,25^. Therefore, we also probed the effect of MLCK-driven contractility on cell identity by expressing a constitutively active mutant. Consistent with the effects of ROCK2^CA^, MLCK^CA^-positive clusters failed to commit to the primitive streak and mesoderm lineage and to trigger the EMT program required for mesoderm identity (**Supplementary Figure 1I-K**). These data support the surprising conclusion that activation of ROCK/MLCK-mediated contractility is sufficient to counteract the effect of WNT signaling activated by CHIR and completely prevents mesoderm conversion and EMT.

### Genetic suppression of actomyosin contractility promotes stem cell conversion to the mesoderm lineage

This unexpected finding prompted us to investigate whether inhibition of contractility might conversely promote mesoderm differentiation. We designed a genetic approach to suppress contractility in human cells. Dephosphorylation of ppMLC is mediated by MLC Phosphatase, a heterotrimer composed of a catalytic subunit PP1cβ, a Myosin phosphatase-targeting regulatory subunit (MYPT1) and a protein of unknown function M20 ^26,27^ (**Figure 2A-B**). Based on previous work describing an optogenetic construct to control actomyosin contractility ^28^, we created a truncated MYPT1, fused with a nuclear export sequence and a mNeonGreen (mNG) reporter (**Figure 2B**), which is expected to act as a constitutively active mutant and promote MLC2 dephosphorylation.

**Figure 2:**
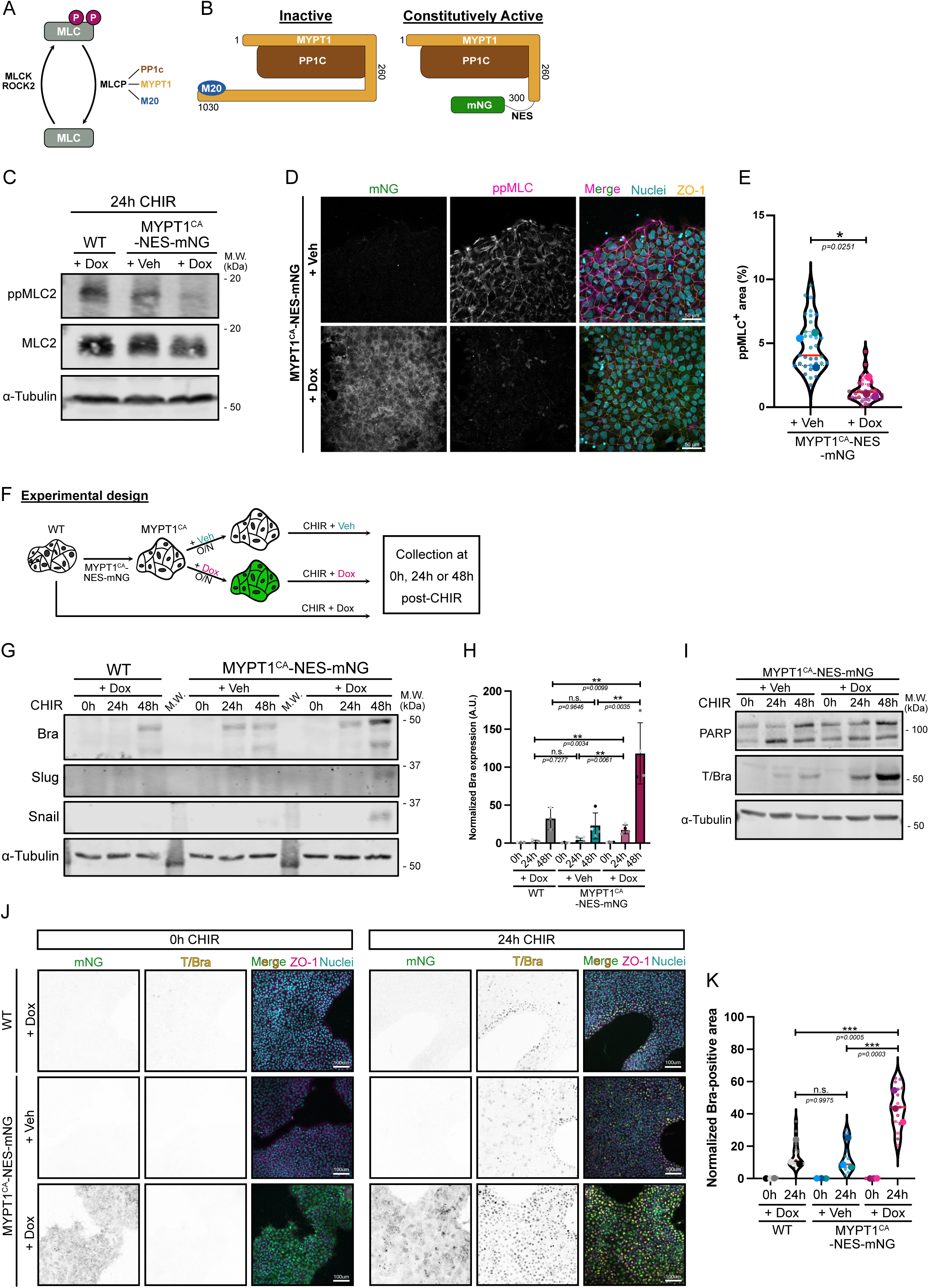
Genetic inhibition of actomyosin contractility promotes hiPSC conversion to the mesoderm lineage. (A) Summary of the main regulators of MLC phosphorylation. (B) Schematic representation of full length human MYPT1 (1030 aa), interacting with PP1c phosphatase and a protein of unknown function M20. Truncation of MYPT1 Cterm Δ301-1030 leads to a constitutive recruitment and activation of PP1C. Truncated MYPT1 was fused with NES-mNeonGreen and is referred to as MYPT1^CA^-NES-mNG. (C) Immunoblot of phospho T18/S19 MLC2 (ppMLC2) and total MLC2 in WT hiPSCs treated with Doxycycline or hiPSC expressing MYPT1^CA^-NES-mNG in the presence or absence of Doxycycline. α-Tubulin was used as loading control. Molecular weights (M.W.) are displayed on the right side. (D, E) – Representative MaxIP immunofluorescence of hiPSC expressing MYPT1^CA^-NES-mNG and treated or not with Doxycycline. Cells were fixed and stained for phospho T18/S19 MLC2 (ppML2 - Magenta), DNA (blue) and tight junction marker ZO-1 (Orange). Scale bar = 50 µm (D). Fraction of the cellular area positive for ppMLC2 following Doxycycline induction of MYPT1^CA^-NES-mNG hiPSC and reported as violin plots. Median (Plain red line) and quartiles (Dotted black lines). n=32 technical repeats across N=3 independent biological repeats. Two-tailed unpaired t test was performed on the biological repeats (E). (F) Experimental design. hiPSC cells were transduced with pInducer20-MYPT1^CA^-NES-mNG and stable population was selected with puromycin. MYPT1^CA^ cells were treated or not with doxycycline for 16h and treated with CHIR supplemented or not with doxycycline. Parental WT line was differentiated with Doxycycline. (G, H) Representative immunoblot for Brachyury (mesoderm marker), Snail and Slug (EMT markers) using WT and MYPT1^CA^-NES-mNG hiPSCs, following CHIR treatment (0h-48h) in the presence or absence of Doxycycline, as shown in (F). Molecular weights (M.W.) are displayed on the right side (G). Brachyury expression was quantified by densitometry and normalized to α-Tubulin as loading control across N=3-4 independent biological repeats. Mean and S.D. are displayed. A one-way ANOVA with Tukey’s multiple comparisons post-test was performed (H). (I) Immunoblot of PARP (cell death marker) and Brachyury (Mesoderm marker) during mesoderm commitment (0hr-48hrs CHIR) using MYPT1^CA^-NES-mNG hiPSCs induced +/ Doxycycline, as shown in (F). Molecular weights (M.W.) are displayed on the right side. (J, K) Representative MaxIP immunofluorescences of WT and MYPT1^CA^-NES-mNG hiPSCs treated with CHIR for 24h +/- Doxycycline as shown in (F). Cells were stained for nuclei (DNA), EMT marker (ZO-1 – Inverted LUT) and Mesoderm marker (T/Bra – Inverted LUT). Scale bar = 100 µm (J). Quantification of Brachyury expression is presented as violin plots. Median (Plain red line) and quartiles (Dotted black lines). n=15-16 technical repeats across N=3 independent biological repeats. One-way ANOVA with Šidák’s multiple comparisons post-test was performed on biological repeats (K).

To validate the activity of this new tool, we cloned the construct (referred to as MYPT1^CA^-NES-mNG) under a doxycycline-inducible promoter and established a stable hiPSC line. Global reduction of ppMLC2 was observed by immunoblot and immunofluorescence after doxycycline addition (**Figure 2C-E**). Strikingly, MYPT1^CA^ cells pre-treated overnight with doxycycline and followed by differentiation (**Figure 2F**), displayed enhanced Brachyury expression (**Fig 2G-K**) and earlier induction of the EMT genes Snail and Slug (**Figure 2G**) following Doxycycline induction, compared to controls. Because apoptosis is crucial to enable mesoderm fate specification ^15^ we probed for PARP cleavage as an apoptosis marker but did not detect any difference (**Figure 2I**). We conclude that reducing MLC2-driven contractility significantly accelerates mesoderm specification.

### Promotion of hiPSC and hESC conversion to the mesoderm lineage by actomyosin relaxation is time-dependent

We next used small molecule inhibitors as an orthogonal method to validate our previous findings. This approach allowed us to expand our investigation to hESCs, while also providing better control over the timing of inhibition.

First, hiPSCs were pre-treated overnight with the ROCK inhibitor, Y-27632 to suppress actomyosin contractility at the pluripotent state then differentiated by addition of CHIR in the continued presence Y-27632 (**Figure 3A**). We confirmed that MLC2 phosphorylation was reduced by ROCK inhibition **(Supplementary Figure 2A-B)**. Similar to the genetic approach, pharmaceutical blockage of contractility strongly promoted Brachyury expression **(Figure 3B-C)**. Expression of other key mesoderm (*MESP1* and *TBX6*) and EMT (*SNAI1* and *SNAI2*) genes was also upregulated following Y-27632 treatment **(Figure 3D-G)**. To rule out an effect of ROCK inhibition on global transcription, we examined expression levels of genes unrelated to mesodermal fate and did not detect any response to decreased contractility, suggesting a specific role for actomyosin contractility in mesoderm specification **(Supplementary Figure 2C)**.

**Figure 3:**
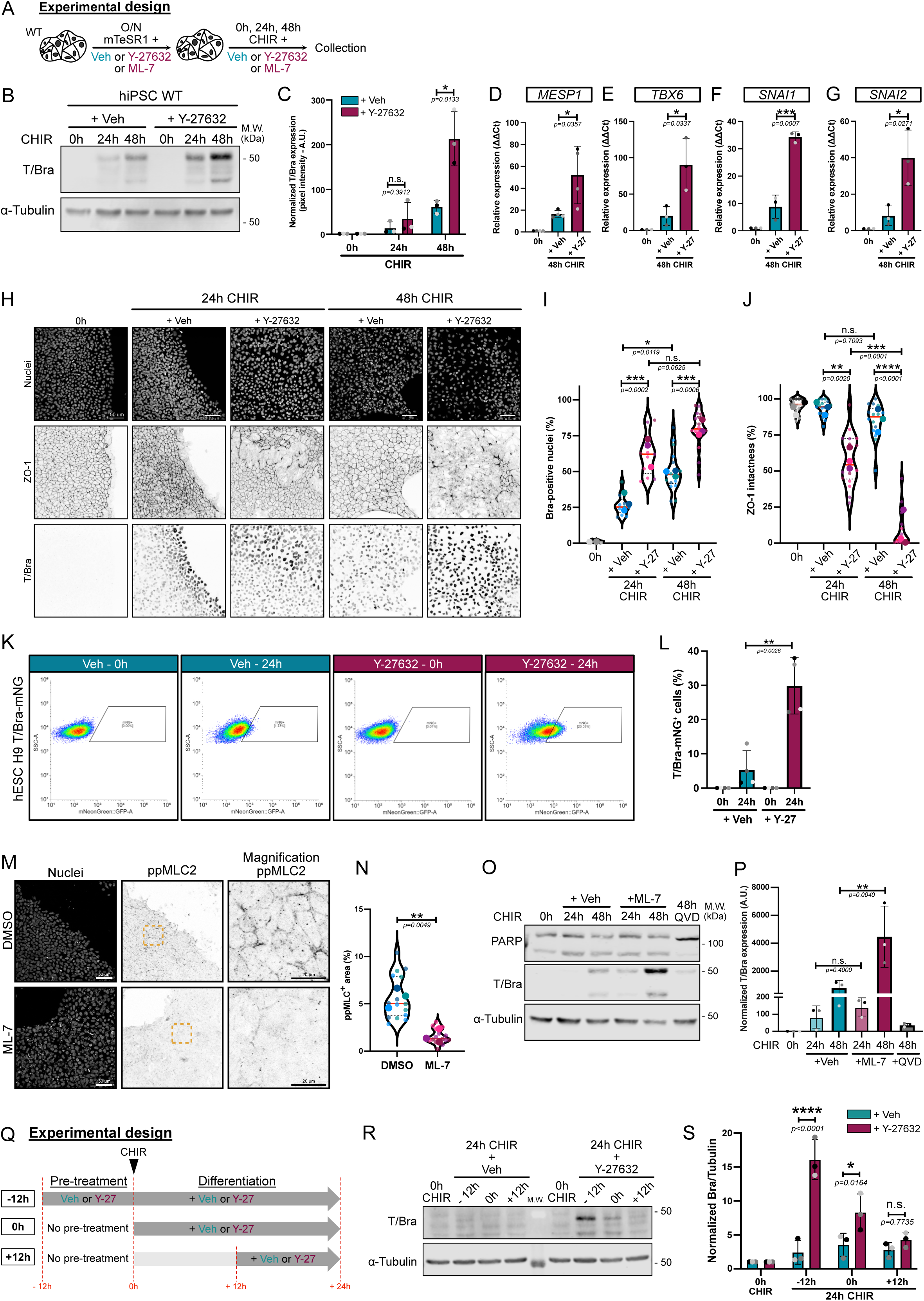
Pharmaceutical inhibition of actomyosin contractility promotes hiPSC and hESC conversion to the mesoderm lineage. (A) Experimental design. WT hiPSCs or hESCs were pre-treated overnight with small molecules to lower contractility, followed by differentiation. (B) Immunoblot of Brachyury during hiPSC-to-mesoderm commitment (0-48h) in the presence (+ Y-27632) or absence (+ Veh) of ROCK inhibitor, as shown in (A). Molecular weights (M.W.) are displayed on the right side. (C) Brachyury expression was quantified by densitometry and normalized to α-Tubulin as loading control across N=3 independent biological repeats. Mean and S.D. are displayed. Two-tailed unpaired t test was performed. (D - G) Relative expression of mesoderm markers (*MESP1, TBX6*) (D, E) and EMT markers (*SNAI1, SNAI2*) (F, G) 48h post CHIR treatment in the presence (+Y-27) or absence (+Veh) of ROCK inhibitor as shown in (A). N=3 independent biological repeats except for *MESP1* N=4 independent biological repeats. Mean and S.D. are displayed. Two-tailed unpaired t test was performed. (H) – Representative MaxIP immunofluorescences during hiPSC-to-mesoderm commitment (0-48h) in the presence (+ Y-27632) or absence (+ Veh) of ROCK inhibitor as shown in (A). Cells were stained with a nuclear marker (DNA), EMT marker (ZO-1 – Inverted LUT) and Mesoderm marker (T/Bra– Inverted LUT). Scale bar = 50 µm. (I, J) Quantification of Brachyury expression (I) and ZO-1 intactness (J) is represented as violin plots with individual measurements (small dots) averaged for each biological repeat (large dots). Median (Plain red line) and quartiles (Dotted black lines). n=10-15 technical repeats across N=3 independent biological repeats. One-way ANOVA with Tukey’s multiple comparisons post-test was performed on biological repeats. (K, L) Analysis of Brachyury expression by flow cytometry following 24h CHIR treatment in the presence (+Y-27632) or absence (+Veh) of ROCK inhibitor, as shown in (A), using Bra-mNeonGreen knock-in H9 hESC line (K). Quantification of the percentage of mNeonGreen-positive cells from N=4 independent biological repeats. Mean and S.D. are displayed. Two-tailed unpaired t test was performed (L). (M, N) Representative MaxIP immunofluorescence of hiPSCs treated with a MLCK inhibitor (ML-7) or Vehicle (DMSO), as shown in (A). Cells were stained for nuclei (DNA) and phospho T18/S19 Myosin Light Chain 2 (ppMLC2 – Inverted LUT). Scale bar = 50 µm. Magnified views of the yellow dotted ROI are shown for ppMLC2. Scale bar = 20 µm (M). Percentage of cellular area positive for ppMLC2 following Vehicle or ML-7 treatment is reported as violin pots. Median (Plain red line) and quartiles (Dotted black lines). n=15 technical repeats across N=3 independent biological repeats. Two-tailed unpaired t test was performed on the biological repeats (N). (O, P) Immunoblot of Brachyury (mesoderm) and PARP (cell death marker) using hiPSCs in the presence (+ ML-7) or absence (+ Veh) of MLCK inhibitor at basal state (0h) and during CHIR treatment (24hrs, 48hrs), as shown in (A). Cells treated with CHIR and Q-VD (Caspase inhibitor) for 48hrs were used as control for PARP immunoblot. Molecular weights (M.W.) are displayed on the right side (O). Brachyury expression was quantified by densitometry and normalized to α-Tubulin as loading control across N=3 independent biological repeats. Mean and S.D. are displayed. Mann-Whitney test was performed for the 24hrs timepoints and two-tailed unpaired t test was performed for the 48hrs timepoints (P). (Q) Experimental design. Effects of Y-27632 was tested at different time along the mesoderm commitment. ROCK inhibitor was added 12h before differentiation (−12h), at the time of differentiation (0h) or 12h after differentiation (+12h). All conditions were collected 24h post differentiation. (R, S) Representative immunoblot for Brachyury following addition of Vehicle or Y-27632 as shown in (Q). Molecular weights (M.W.) are displayed on the right side (R). Brachyury expression was quantified by densitometry and normalized to α-Tubulin as loading control across N=3 independent biological repeats. Mean and S.D. are displayed. Two-way ANOVA with Šidák’s multiple comparisons post-test was performed (S).

At the cellular level, Brachyury expression is initially restricted to the colony edges at 24hrs, before broader expression across the colony at later timepoints (**Figure 3H-I**). However, Y-27632 treatment resulted in more widespread Brachyury expression **(Figure 1H-I)**, markedly accelerated EMT as shown by loss of the epithelial marker ZO-1 from tight junctions (**Figure 1H, J**), and increased expression of the mesenchymal marker Slug **(Supplementary Figure 2D)**. To validate these data, we used a more specific ROCK inhibitor, H-1152, which led to similar decreases in ppMLC2 **(Supplementary Figure 2E-F)** and recapitulated the early EMT and higher induction of mesoderm markers **(Supplementary Figure 2G-M)**. Apoptosis was not affected following Y-27632 or H-1152 treatment **(Supplementary Figure 2L)**. Moreover, these effects of ROCK inhibition were conserved in hESCs expressing a knock-in T-mNeonGreen fusion ^16^ **(Figure 3K-L)** and in wild-type H9 hESCs treated with CHIR (**Supplementary Figure 2N-O**). Importantly, similar to hiPSCs, there was no effect of reduced contractility on apoptosis initiation (**Supplementary Figure 2N and 2P**) or on exit from the pluripotency state **(Supplementary Figure 2N and 2Q)**.

Because MLCK activation was sufficient to block cell commitment (**Supplementary Figure 1I-K**), we tested the effect of ML-7, a potent MLCK inhibitor ^29^. ML-7 led to efficient reduction in ppMLC2 **(Figure 3M-N)** and strongly enhanced Brachyury expression at 48hrs, without affecting cell death **(Figure 3O-P)**. Together these data show, unexpectedly, that pharmacological blockade of MLC2-mediated force generation enhances mesoderm commitment quantitatively and accelerates the EMT required for conversion to cardiac mesoderm.

These results were particularly surprising, because hiPSC colonies show rapid retraction of their edges following CHIR addition **(Supplementary Figure 3A and Supplementary Video 1)**, consistent with observations from other groups ^30^. Having previously reported an essential and permissive role for apoptosis during mesoderm specification ^15^, we tested whether colony retraction was related to the wave of cell death that follows CHIR addition. However, co-treatment with a pan-Caspase inhibitor (Q-VD-OPH) did not prevent retraction of colony edges **(Supplementary Figure 3A)**, ruling out an effect of cell density and cell death.

Commitment to the mesoderm lineage also resulted in thicker epithelial layers, containing taller cells **(Supplementary Figure 3B-D)**, suggesting morphological changes that are highly dependent on the cellular contractile state. Consistent with the role of nuclei as a mechanosensor ^31,32^, differentiating cells exhibit significantly smaller nuclear area after 48h of CHIR treatment **(Supplementary Figure 3E-F)**. To address the variation of cell density, we correlated each of our nuclear area measurements with the actual cell number **(Supplementary Figure 3G)**. As expected, nuclear area negatively correlates with cell density before differentiation. However, we did not find any correlation following CHIR treatment, together suggesting that the smaller nuclear size at 48h of differentiation is unlikely to be related to cell density.

While these observations were indirect, they point towards a change in contractile status during mesoderm commitment. To directly test for higher contractility, differentiating cells were stained ppMLC2. Immunofluorescence revealed a gradual increase of ppMLC2 intensity over time **(Supplementary Figure 3H-I)**. Finally, we took advantage of the birefringence property of actin fibers to measure optical retardance using quantitative polarization microscopy ^33^. Birefringence of a material changes proportionally to the applied strain, providing an orthogonal way to probe for contractile status during mesoderm commitment. Confirming our previous data, retardance values increased during hiPSC-to-mesoderm conversion **(Supplementary Figure 3J)**. Specificity of the retardance value as a readout for contractility was tested by incubating undifferentiated hiPSCs with Y-27632, which reduced retardance. Together, these data suggest that hiPSC differentiating along the cardiac mesoderm trajectory, in the absence of any external perturbation, experience increasing contractile forces mediated by MLC2 phosphorylation.

How can we reconcile the surprising observation that differentiating cells intrinsically become more contractile, with the inhibitory function of actomyosin contractility on mesodermal specification? We wondered if the timing of inhibitor treatment was important, and therefore staggered the timing such that Y-27632 was added either 12 hrs prior differentiation (−12h), at the time of differentiation (0h) or 12 hrs following CHIR treatment (+12h), and analyzed Brachyury expression after 24hrs post-CHIR addition (**Figure 3Q**). Pre-treatment of pluripotent stem cells with ROCK inhibitor (−12h) strongly promoted mesoderm differentiation. Co-treatment with ROCK inhibitor and CHIR reduced the effect, and addition of ROCK inhibitor at +12hrs had no effect on differentiation (**Figure 3R-S**). We conclude that reduced contractility primes pluripotent stem cells for specification but has no effect once specification has been initiated by WNT activation.

### Hippo pathway and force-dependent WNT ligand secretion do not contribute to enhanced mesoderm specification

Mechanistically, how do changes in contractility impact cell fate specification? A major target of mechanical force in epithelia is the Hippo pathway. YAP, a downstream effector of Hippo signaling, has been previously associated with cell fate patterning in 2D gastruloids ^34^, and substrate stiffness-driven mesoderm specification ^35^. Low tension activates this pathway, resulting in YAP phosphorylation and degradation. Conversely, high tension suppresses Hippo signaling, blocking phosphorylation of YAP. The non-phosphorylated YAP translocates to the nucleus to regulate gene expression. Surprisingly, however, phosphorylated YAP levels were not affected during hiPSC differentiation or by cell relaxation (**Supplementary Figure 4A, B**). We next directly probed for *CTGF* and *CYR61* expression, two well-described YAP target genes. Gene expression strongly decreased during differentiation, as expected given the reported repressive functions of YAP during mesendoderm specification ^36^, but expression level was not affected by the cellular contractile status (**Supplementary Figure 4C**). Together, these data rule out a significant role for the Hippo pathway in the acceleration of mesodermal differentiation following cell relaxation.

Next, we investigated the positive feedback loop reported for BMP4-driven mesoderm commitment, which relies on tension-dependent secretion of canonical WNT ligands ^16^. We treated our MYPT1^CA^-NES-mNG cells with Doxycycline to trigger cell relaxation and probed for canonical and non-canonical WNT ligands. As expected, expression of *TBX6* (a mesoderm marker) increased in low contractile cells. Interestingly, canonical WNT ligand expression (*WNT3A* and *WNT8A*) was strongly upregulated by suppression of contractility, while non-canonical *WNT4* expression was unaffected by CHIR treatment or by relaxation (**Supplementary Figure 4D**). To test whether increased *WNT3A/8A* expression was responsible for increased mesoderm commitment in low-contractility cells, we blocked WNT ligand processing and secretion using IWP-2, a Porcupine inhibitor. Addition of IWP-2 did not reverse mesoderm gene expression in Y-27632-treated cells, suggesting that this pathway does not drive accelerated mesoderm specification in response to cell relaxation (**Supplementary Figure 4E**). Together, these data ruled out the involvement of two major pathways, previously thought to link cell tension to mesoderm commitment ^16,35^.

### Intercellular adhesion and Adherens Junctions mediate cell contractility

Seeking a potential mechanism, we turned our attention to intercellular adherens junctions (AJs). These junctions are under tension, generated by actomyosin interactions with vinculin/α-catenin, which in turn are coupled to β−catenin and E-cadherin ^37^. In addition, tension across AJs is mediated by homotypic and calcium-dependent interactions between E-cadherin molecules on adjacent cells ^38^. As a major signaling hub, we reasoned that AJs might transduce mechanical forces to impact cell fate specification. We first tested whether disrupting AJs using EGTA (a calcium chelator) would affect cell specification. EGTA treatment caused colony decompaction (**Figure 4A and Supplementary Fig 5A**), consistent with previous literature linking AJ-disengagement with reduced force transmission ^39,40^. Strikingly, EGTA treatment triggered a strong enhancement of mesodermal and EMT marker expression (**Figure 4B-E and Supplementary Figure 5B-C**). Knowing that reduced contractility mimics this phenotype, we tested whether EGTA treatment would alter contractility. Staining for ppMLC2 confirmed that EGTA treated cells are in a low contractile state (**Supplementary Figure 5D-E**).

**Figure 4:**
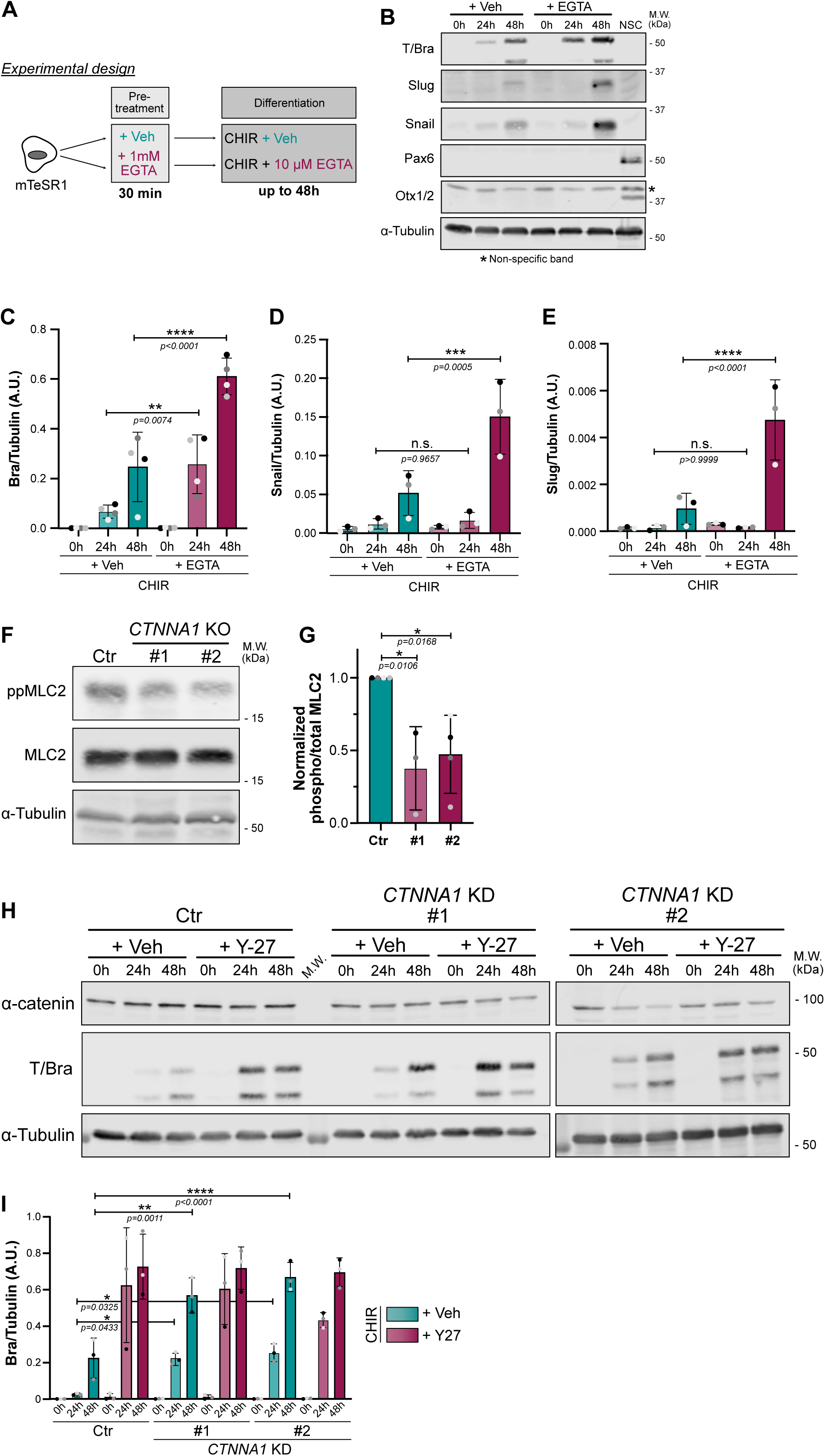
Adherens junctions disengagement promotes mesoderm fate. (A) Experimental design. hiPSCs were pre-treated with an acute concentration of EGTA to disrupt calcium-dependent E-Cadherin junctions. Following pre-treatment, hiPSCs were differentiated in the presence of a lower concentration of EGTA. (B-E) Representative immunoblot of hiPSCs treated as shown in (A) and probed for Brachyury (mesoderm marker), Snail and Slug (EMT markers) and Pax6 and Otx1/2 (Neuroectoderm markers as negative controls). Neural Stem Cell (NSC) lysate was used as positive control for Pax6 and Otx1/2. Non-specific band is designated by an asterisk. Molecular weights (M.W.) are displayed on the right side (B). Expression of Brachyury (C), Snail (D) and Slug (E) was quantified by densitometry and normalized to α-Tubulin across N=3 independent biological repeats. Mean and S.D. are displayed. One-Way ANOVA with Šidák’s multiple comparisons post-test was performed. (F, G) Representative immunoblot of control (Ctr) and α-catenin knockdown hiPSC lines at basal state and probed for total and phospho-MLC2. Molecular weights (M.W.) are displayed on the right side (F). Quantification of active MLC2 (ppMLC2/MLC2) was measured by densitometry across N=3 independent biological repeats. Mean and S.D. are displayed. One-Way ANOVA with Dunnett’s multiple comparison post-test was performed (G). (H, I) Representative immunoblot of control (Ctr) and α-catenin knockdown (*CTNNA1* KD #1 and #2) hiPSC lines, differentiated for up to 48hrs in the presence or absence of ROCK inhibitor Y-27632 and probed for α-catenin, Brachyury and α-Tubulin as loading control. Molecular weights (M.W.) are displayed on the right side (H). Normalized Brachyury expression was measured by densitometry across N=3 independent biological repeats. Mean and S.D. are displayed. One-Way ANOVA with Šidák’s multiple comparisons post-test was performed (I).

As an orthogonal approach, we sought to uncouple force transmission from the actomyosin network to E-cadherin via vinculin/α-catenin. To this end, we created α-catenin knockdown (*CTNNA1* KD) hiPSCs, with a 50% decrease in protein expression (**Supplementary Figure 5F-G**). First, because AJs are required for stemness acquisition during fibroblast reprograming to iPSCs ^41^, we checked that the *CTNNA1* knockdown cells had not lost pluripotency (**Supplementary Figure 5F, H-I**). Notably, ppMLC2 levels were 50% lower in the KD cells (**Figure 4F-G**), confirming that α-catenin is a major contributor to cell actomyosin activity. The *CTNNA1* KD hiPSCs exhibited higher and earlier expression of Brachyury. Strikingly, however, these cells failed to further respond to Y-27632 treatment, while control cells responded as expected by increasing Brachyury expression (**Figure 4H-I**). Together these data demonstrate that AJ disruption reduces actomyosin contractility and that α-catenin-mediated force transmission is a major determinant of mesoderm specification kinetics.

### Junctional β-catenin localization scales with intrinsic actomyosin contractility in undifferentiated hiPSCs

Our discovery of a key role for AJ mechanics in cell fate specification suggested that β-catenin, which in epithelial cells is mostly bound to E-Cadherin at AJs, might respond to changes in actomyosin contractility by modulating WNT signaling responsiveness. Release of β-catenin from AJs, for example, might increase the cytoplasmic and nuclear pools to amplify expression of *TBXT* and other mesoderm genes. Mechanosensitive functions for β-catenin during development and tissue homeostasis have been described ^16,42^ but whether the junction-associated and cytoplasmic β-catenin pools both contribute to WNT signaling is still controversial ^18^, and other studies have suggested a connection in which increased junctional β-catenin correlates with increased nuclear β-catenin ^17^.

We hypothesized that loss of tension at AJs would release β-catenin to promote WNT-responsive gene expression, and conversely that increased tension would promote binding at AJs, thereby titrating out the free cytoplasmic/nuclear β-catenin. To test this mechanism, undifferentiated hiPSCs were treated with either the Rho activator CN03 or ROCK inhibitor Y-27632, causing an increase or decrease in ppMLC2, respectively (**Figure 5A-B and Supplementary Figure 6A**). Strikingly, under each condition, junctional β-catenin levels correlated closely with the level of active ppMLC2 in the cells (**Figure 5B-C and Supplementary Figure 6B**). To rule out staining artifacts, we performed comparable experiments using our EGFP-β-catenin knock-in hiPSC line (**Figure 5D**), with a similar conclusion (**Figure 5E-F, Supplementary Figure 6C and Supplementary Video 2**). Importantly, overall β-catenin expression is not affected by contractile state, suggesting that contractility does not regulate β-catenin level but rather its localization (**Supplementary Figure 6D-F**). Together, these data demonstrate that β-catenin association with AJs scales directly with actomyosin contractility.

**Figure 5:**
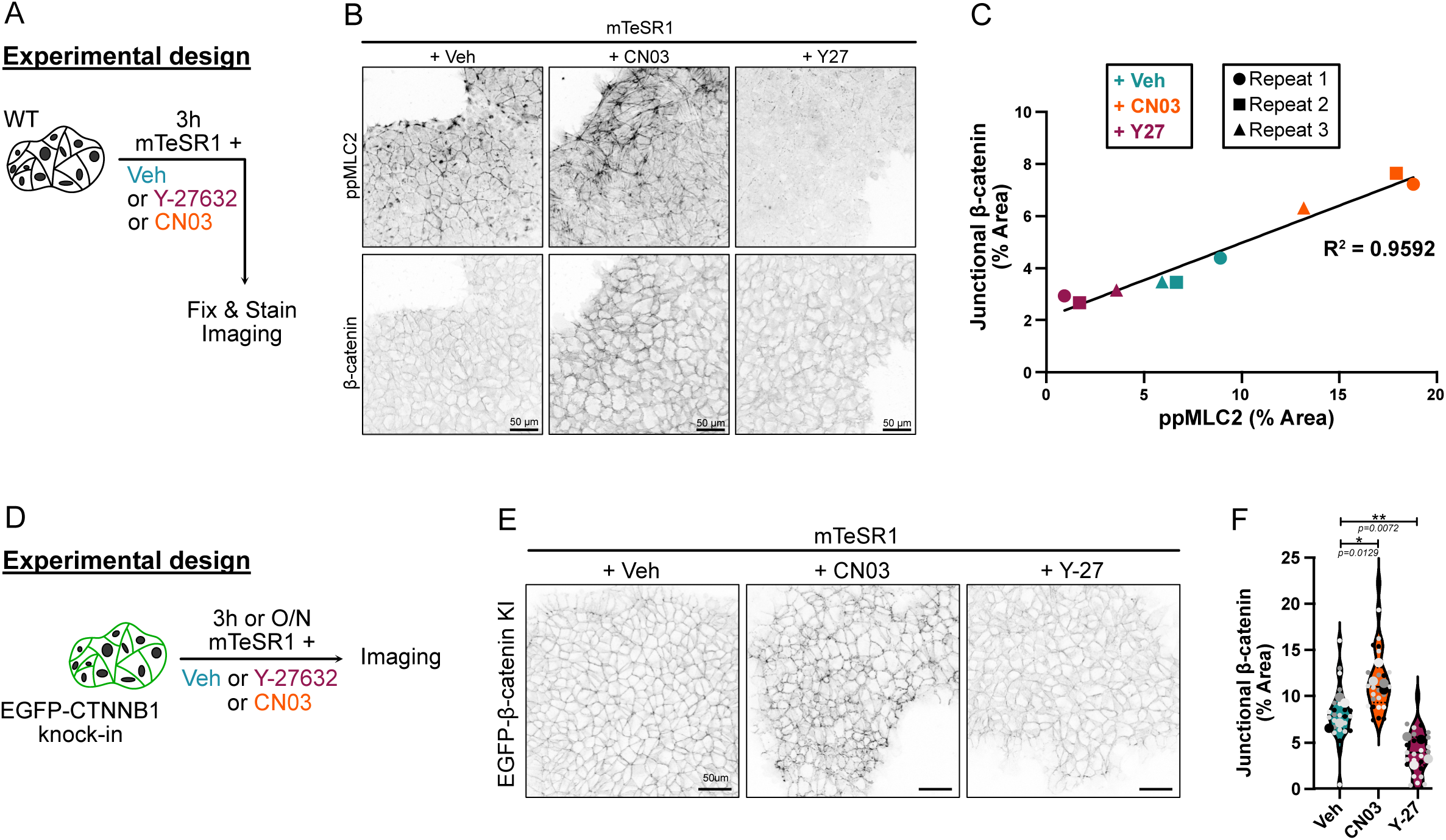
Contractility differentially regulates β-catenin localization at adherens junctions. (A) Experimental design (B) Representative immunofluorescence of fixed hiPSCs treated with Vehicle, 4µg/mL CN03 or 10 µM Y-27632 for 3h and stained for ppMLC2 and β-catenin (inverted LUT), as shown in (A). Scale bare = 50 µm. (C) Linear regression analysis of junctional β-catenin and cellular ppMLC2 was quantified across N=3 independent biological repeats (representing n=16 for Vehicle, n=19 for CN03 and n=17 technical repeats for Y-27632). R^2^ value is reported. (D) Experimental design (E, F) Representative still pictures from mEGFP-β-catenin knock-in (KI) hiPSCs treated with Vehicle, 4µg/mL CN03 or 10 µM Y-27632 for 3h, as shown in (D). Scale bare = 50 µm (E). Quantification of junctional β-catenin is reported for the different treatment across N=4 independent biological repeats (representing n=25 technical repeats). Violin plots representing median and quartiles. One-Way ANOVA with Dunnett’s multiple comparison post-test was performed on biological repeats (F).

### Cell relaxation promotes nuclear β-catenin localization and occupancy at *TBXT* promoter

In the absence of WNT signaling, cytosolic β-catenin is maintained at a low level by association with a destruction complex, where it is phosphorylated at Ser45 by CK1, which primes the protein for subsequent phosphorylation by GSK3β, driving its ubiquitination and degradation. WNT activation disrupts the destruction complex, permitting the accumulation of stable, non-phosphorylated β-catenin, which can enter the nucleus to drive gene expression. This “active” state can be assessed from the detection of non-phosphorylated Ser45 β-catenin. To test for differential accumulation of active β-catenin, we isolated cytosolic and nuclear fractions from cells that contain the Dox-inducible MYPT1^CA^-NES-mNG construct (**Supplementary Figure 7A**). Purity of each fraction was assessed for enrichment of a nuclear and cytosolic marker, Lamin A/C and GAPDH, respectively (**Supplementary Figure 7B**). Fractionation showed that active β-catenin was strongly enriched in both the cytosolic and the nuclear fractions of cells treated with CHIR + Dox (24h) versus cells treated only with CHIR (**Figure 6A-B**). Immunostaining confirmed the nuclear enrichment of active β-catenin (**Figure 6C and Supplementary Figure 7C-D**) after 24h of differentiation in low contractile conditions.

**Figure 6:**
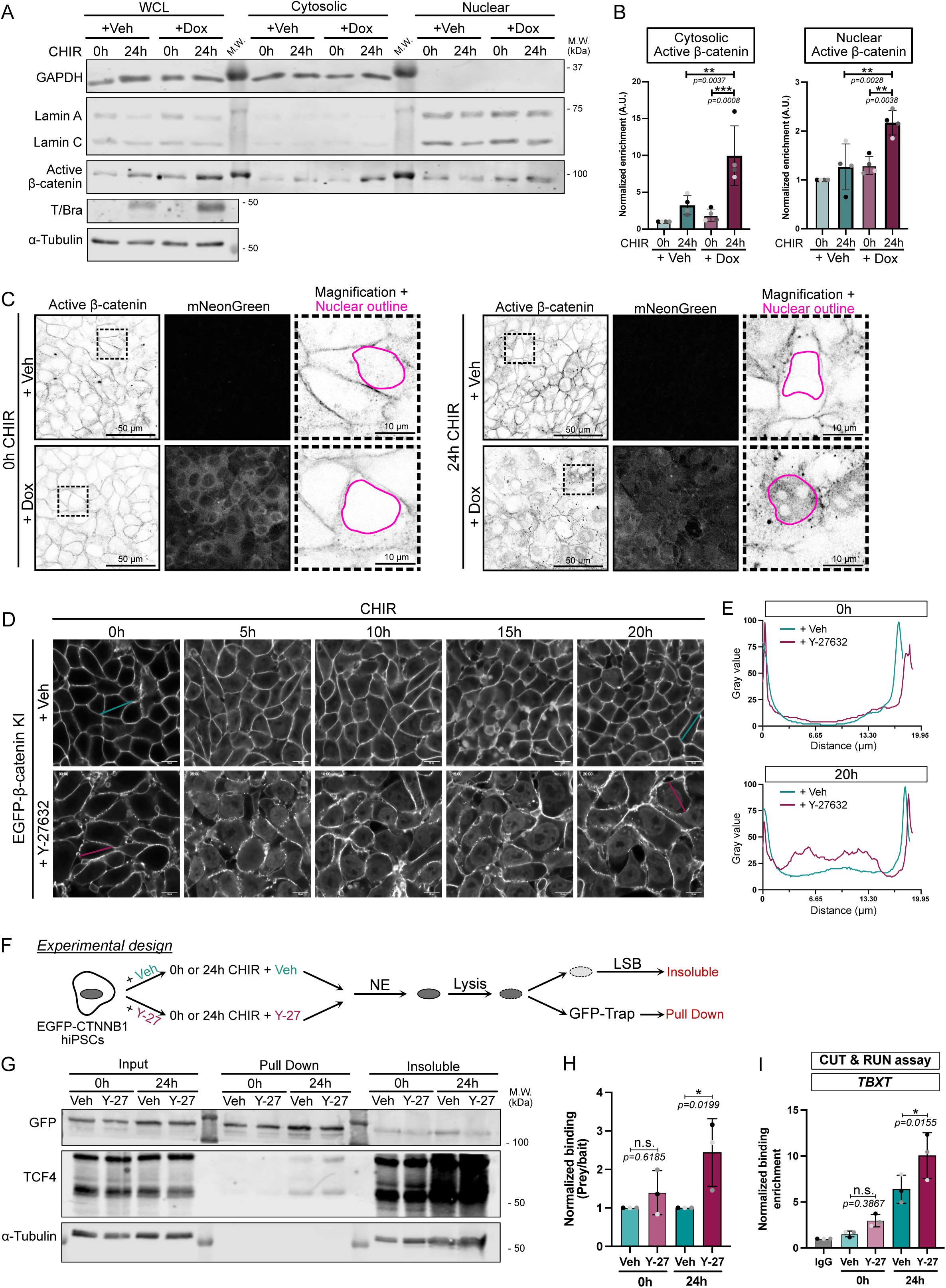
Reduced cell contractility promotes β-catenin nuclear activity and enhances binding to mesodermal gene. (A) Representative immunoblot as explained in Supplementary Figure 6A. MYPT1^CA^-NES-mNG hiPSCs were induced with CHIR for 24hrs in the presence or absence of Doxycycline. Whole cell lysate (WCL) was collected and loaded as input while cytosolic and nuclear fractions were extracted. Fraction purity was probed using cytosolic (GAPDH) and nuclear (Lamin A/C) markers. Active β-catenin (non-phospho S45) enrichment was probed for each fraction and α-Tubulin was used as loading control. Molecular weights (M.W.) are displayed on the right side. (B) Quantification of active β-catenin enrichment in the cytosolic (left) and nuclear (right) fractions. N=4 independent biological repeats. Mean and S.D. are displayed. One-way ANOVA with Šidák’s multiple comparisons post-test was performed. (C) Representative immunofluorescence for active non-phospho S45 β-catenin (Inverted LUT – confocal slice) and mNeonGreen (MaxIP). MYPT1^CA^-NES-mNG hiPSCs were fixed at 0h and 24h post CHIR treatment +/- Doxycycline. Scale bar = 50 µm. Insets represent a magnified view of the active non-phospho S45 β-catenin with an example of the nuclei binary mask overlaid (pink). Scale bar = 10 µm. (D, E) Stills confocal timelapse imaging of mEGFP-β-catenin knock-in hiPSCs treated with CHIR +/- Y-26732 (Related to Supplementary movie 3). Time is shown in hours. Scale bar = 10 μm (D). mEGFP-β-catenin intensity is reported as a scan line plot across a cell at 0h and 20h (E). (F) Experiment design for GFP-Trap immunoprecipitation. mEGFP-β-catenin knock-in hiPSCs were pre-treated with Vehicle or Y-27632 and differentiated for 24h. Nuclei were isolated and nuclear lysates were mixed with GFP-Trap beads. (G, H) Representative GFP-Trap immunoprecipitation of mEGFP-β-catenin knock-in hiPSCs, probed for TCF7L2/TCF4 and α-Tubulin as loading control. Molecular weights (M.W.) are displayed on the right side (G). Quantification of TCF4 (Prey) binding to mEGFP-β-catenin (Bait) was measured by densitometry across N=3 independent biological repeats. Mean and S.D. are displayed. One-way ANOVA with Šidák’s multiple comparisons post-test was performed. (I) CUT & RUN assay for IgG or β-catenin binding to *TBXT* promoter in WT hiPSC pre-treated with Veh or Y-27632 (0h CHIR) or following 24h of CHIR treatment in the presence of absence of ROCK inhibitor. N=3 independent biological repeats. Mean and S.D. are displayed. One-way ANOVA with Šidák’s multiple comparisons post-test was performed.

We validated this observation by imaging endogenous mEGFP-β-catenin knock-in hiPSCs treated with CHIR in either control or low contractile (+Y-27632) conditions. First, we noticed nuclear accumulation of mEGFP-β-catenin within 5h of CHIR treatment, regardless of the contractile status of the cells. However, while β-catenin signal decreased over time in vehicle-treated cells, remaining visible after 20h in only a few nuclei, low contractile cells showed a robust and enhanced nuclear localization (**Figure 6D-E and Supplementary Video 3**).

Finally, knowing that β-catenin is enriched in the nucleus upon low contractility, we focused on the ability of β-catenin to interact with its binding partners and to regulate WNT-responsive genes. Because of the timing of our phenotype, we ruled out an involvement of β-catenin interactions with transcription factors (such as SOX family, SMAD, TBX3) ^43,44^, as they are involved later during specification ^45^. In addition, ChIP-seq experiments report that TCFs/β-catenin binding is the main event during pluripotency exit and early specification ^45,46^.

Mammalian genomes encode four TCF proteins (TCF7, TCF7L1, TCF7L2 and LEF1), all of which bind to similar DNA sequences ^47^, show redundant functions ^46,48^ and co-occupy similar DNA regions ^49,50^. TCF3 (encoded by *TCF7L1*) is the most highly expressed member in hiPSC (**Supplementary Figure 7E**) but is widely reported to be repressive ^51–54^. Therefore, we focused on the second most highly expressed factor, TCF4 (encoded by *TCF4L2*). Nuclei isolated from Y-27632-treated EGFP-β-catenin knock-in hiPSCs showed higher binding between β-catenin and TCF4 compared to vehicle-treated cells (**Figure 6F-H**). Finally, CUT & RUN assays showed that low contractility significantly increased β-catenin recruitment to *TBXT* promoter during differentiation (**Figure 6I and Supplementary Figure 7F**), supporting a priming model. Together, these findings suggest that the promotion of nuclear β-catenin accumulation by cytoskeletal relaxation accelerates mesoderm program by binding to its TCF co-mediator. However, this model can only hold true if the concentration of CHIR used in our study (7.5 µM) does not saturate the β-catenin signaling response. To test this premise, we doubled the concentration of CHIR and probed for expression of WNT target genes as a read-out for WNT signaling (**Supplementary Figure 7G**). *LEF1* and *AXIN2* expression responded in a dose-dependent manner to CHIR. Interestingly, however, at any given CHIR concentration, inhibition of cell contractility further enhanced WNT-target gene responses.

Collectively, we demonstrate that actomyosin contractility regulates β-catenin availability and that cell relaxation amplifies active β-catenin translocation to the nucleus synergistically with CHIR to enhance mesoderm commitment (**Figure 7**).

**Figure 7:**
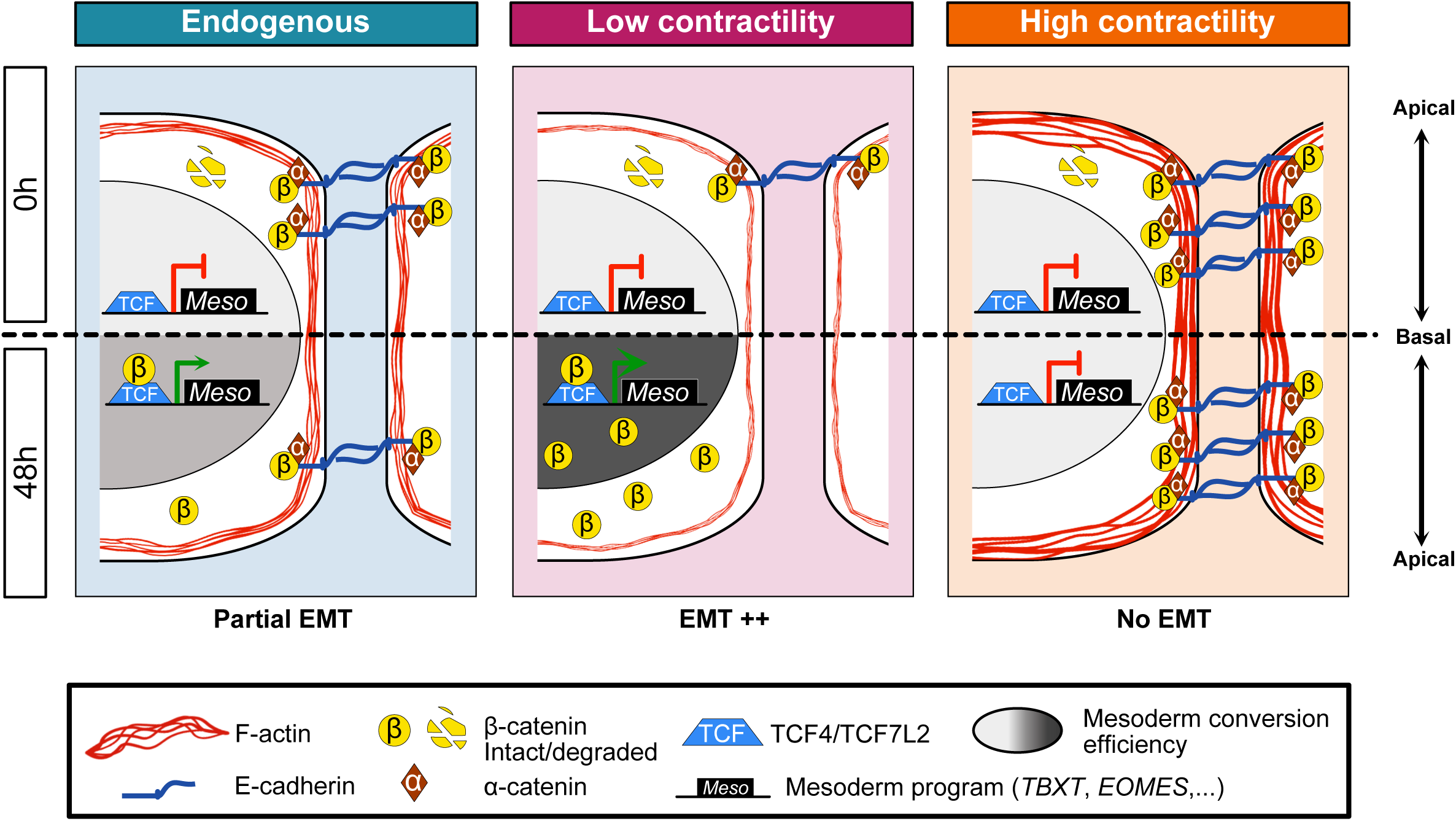
Working model. Effect of actomyosin contractility on mesoderm lineage commitment. Each panel represents a specific contractile status at basal state (0h - top) and after 48h (bottom) of CHIR treatment. At basal state, β-catenin is degraded and localized at AJs. The amount of junctional β-catenin correlates with the level of contractility experienced by hiPSCs. During differentiation, β-catenin accumulates in the nucleus and activates a mesoderm program through its binding with TCF4.

## Discussion

Here, we have identified an inverse relationship between actomyosin contractility and mesoderm specification in hPSCs. Mechanistically, our data support a model in which adherens junctions serve as dynamic reservoirs that sequester β-catenin away from the cytoplasmic signaling pool. We propose that the inter-dependence of actomyosin contractility and AJs creates a titration mechanism for β-catenin availability, thereby tuning mesoderm specification. Previous papers have reported that, contrary to our observations, increased mechanical tension promotes β-catenin-dependent mesoderm induction. For example, tension has been reported to promote Brachyury expression in hESC ^17^, in Bilateria ^55^ and Cnidaria ^20^, while in gastrulation-like models of hESC, Muncie et al argued that high cell adhesion tension on compliant substrate can promote mesoderm induction ^16^. This study proposed a dual function for mechanical tension. First, BMP4-driven Brachyury expression relies on mechanical stretch of cell junctions, allowing destabilization of β-catenin from AJs and its nuclear translocation. Following this initiation stage, a positive feedback loop reinforces mesoderm commitment through increased WNT ligand secretion. Strikingly, on Matrigel-coated plates, we observe the opposite outcome. Importantly, our unconstrained system directly focuses on contractility rather than tension geometry or substrate properties. Our findings suggest that intrinsic contractility, even in unpatterned settings, is sufficient to guide cell fate decisions. We provide causal and cell-intrinsic evidence that actomyosin-generated forces act upstream of mesodermal fate, in the pluripotent state, to inhibit cell specification. Second, our protocol directly activates WNT signaling by inhibition of the destruction complex, as do WNT ligands, in contrast with the indirect effects of BMP4 activation. Importantly, a previous study showed that epithelial cohesiveness and tight junctions block access by BMP4 to its receptor on the lateral plasma membrane ^56^. Therefore, BMP4 signaling can only activate Brachyury expression at colony edges and corners ^16^. However, our protocol uses a cell-permeable small molecule inhibitor as the differentiating cue, circumventing this diffusion barrier. Third, our findings distinguish our model from alternative force-mediated mechanisms such as Hippo/YAP signaling ^35^ or tension driven WNT ligand secretion ^16^, neither of which accounted for the enhanced mesoderm specification. One explanation might be that pluripotent stem cells are insensitive to Hippo activation because of their inability to perceive cell-cell interactions, as previously reported ^35^. Finally, our system relies on activating or reducing contractility at the pluripotent state by pre-treating cells prior to differentiation and maintaining treatment during cell commitment. We observed a timing-dependent response during which enhanced mesoderm identity was only achieved following ROCK inhibition pre-treatment of cells in the pluripotent state. Therefore, our study suggests that low contractility primes pluripotent cells to enter the differentiation trajectory. Such a licensing mechanism would likely involve cis-regulatory element-mediated chromatin priming that can potentiate gene activity prior to specification events ^57^. Alternatively, we note that actomyosin-dependent epigenome regulation was previously associated with RNA-polymerase II activity and lineage commitment in adult stem cells ^58^.

We here describe a titration mechanism by actomyosin contractility that allows a synchronization with developmental processes during cell commitment. β-catenin exists in two pools, one bound to the adherens junctions through E-cadherin and one cytosolic pool that is central to WNT signaling. The regulation and exchange of β-catenin molecules between these two pools remains poorly understood. E-cadherin binding to β-catenin prevents its nuclear translocation and transcriptional co-activator function ^19^. Src-mediated phosphorylation of β-catenin at residue Y654 has been reported to reduce binding to E-cadherin ^59^, so one possibility might be that Src tyrosine kinase activity is anti-dependent on contractility in pluripotent stem cells. Alternatively, however, Gayrard et al ^21^ reported an inverse mechanism through which, in migrating cells, Src-dependent phosphorylation of FAK causes actin remodeling and E-cadherin tension relaxation, resulting in the release and nuclear translocation of β-catenin. In our system, reduced tension causes loss of β-catenin from AJs, accumulation in the cytosolic compartment, and translocation into the nucleus. Finally, in *Xenopus*, decreased ROCK-dependent actomyosin contractility promotes an ectoderm-to-mesoderm transition, suggesting force-dependent lineage plasticity is a conserved mechanism ^60^. While we have not tested for alternative cell fate following CHIR addition, our differentiation protocol was adapted from Lian et al ^12,13^ and was not reported to lead to ectodermal identity.

## Acknowledgements

We thank Cynthia Reinhart-King for giving us access to the QPOL facility. We thank the Weaver lab (UCSF) and the Gardel lab (University of Chicago) for sharing the T-mNG knock-in hESC line. We thank Dr. Vivian Gama lab (Vanderbilt University) for providing the NSC lysate, Pax6 and Otx1/2 antibodies. We also thank all members of the Macara lab, especially Dr. Christian de Caestecker for valuable suggestions and advice. This work was funded by an American Heart Association postdoctoral fellowship (Award ID 23POST1018347 to L.F.), a 2024 Vanderbilt Mechanobiology fellowship (to L.F.), a 2024 and 2025 Vanderbilt Mechanobiology fellowship (to L.F.) and a grant from the National Institutes of Health (GM070902 to I.G.M.).

## Author contributions

L.F. conceived the study, performed and quantified all experiments except the ones stated below, and wrote the manuscript. V.V. performed the CUT & RUN assay. W.W. imaged and quantified the QPOL data. I.G.M. supervised the project, and edited the manuscript.

## Data availability statement

The data are available from the corresponding author upon reasonable request.

## Material & Methods

### Research compliance and ethical regulation

All experiments using human ESCs were performed using the WA09 (H9) cell line under the supervision of the Vanderbilt Institutional Human Pluripotent Cell Research Oversight Committee (protocol IRB no. 160146). No cells were sourced directly from human embryos.

### Reagents

Common laboratory reagents used in this study are listed in Supplementary Table 1

### Cell culture and maintenance

Human GM25256 iPSCs were obtained from the Coriell Institute. The mEGFP–CTNNB1 knock-in GM25256 iPSC cell line was obtained from the Allen Cell Collection, Coriell Institute (cell ID AICS-0058-067). The human ESC line H9 (WA09) was obtained from the WiCell Research Institute. The T-mNG knock-in hESC H9 was made by Valerie Weaver lab (UCSF) and kindly provided by Margaret Gardel (University of Chicago). The iPSC and ESC H9 lines were cultured on Matrigel-coated six-well plates (Matrigel diluted at 42 μg ml−1 in DMEM/F12 medium) and cultured in mTeSR1 medium. The medium was changed daily until the cells reached 70% confluency. Cells were passaged using Gentle cell dissociation reagent for 4 min, resuspended in mTseR1 medium as small clusters and replated at 1:7. HEK293T cells were obtained from the ATCC. Cells were grown in 10% FBS DMEM medium and passaged at 1:10 every 2-3 days. All cell lines used in this study were maintained at 37 °C under 5% CO2.

### Cell freezing and thawing

Human iPSCs and ESCs were harvested from culture dishes using Gentle cell dissociation reagent and centrifuged at 120g for 3 min. The pellets were resuspended in mTeSR1 medium supplemented with 10% DMSO and aliquoted into cryovials. Cells were first transferred to −80 °C for 24 h before long-term storage in liquid nitrogen. iPSCs and ESCs were slowly thawed using mTeSR1, centrifuged and resuspended in mTeSR1 supplemented with 10 μM Y-27632 for 24 h.

### Mesoderm differentiation protocol and drug treatments

Differentiation was initiated when cells reach 70-80% confluency. Cells were treated with 7.5 µM CHIR99021 (hereafter CHIR) diluted in RPMI media supplemented with 1X B27 minus insulin and 100U/mL Penicillin and 100 μg/mL streptomycin (1X).

For pharmaceutical inhibition of contractility, cells (50% confluent) were pre-treated overnight or 3h with drug or vehicle diluted in mTeSR1. Differentiation was initiated the day after (cells reached 70-80% confluent) by switching to CHIR media as described above, supplemented with the drug or vehicle. For MYPT1^CA^, cells were pre-induced overnight with Doxycycline or vehicle (water) diluted in mTeSR1 before initiating the differentiation protocol in the presence of Doxycycline or water.

For pharmaceutical stimulation of cell contractility, 70-80% confluent cells were pre-treated for 3h with 4 μg/mL CN03 or water in mTeSR1, followed by CHIR treatment supplemented with CN03 or water. For EGTA treatment, 70-80% confluent cells were pre-treated for 30min with 1mM of EGTA or water in mTeSR1, followed by CHIR treatment supplemented with 10 μM EGTA or water. For concentration of each drug, please refer to Supplementary Table 1.

### Lentiviral preparation

HEK293T were grown at 40% confluency in 10 cm plate and transfected using Calcium-Phosphate. Briefly, 20 μg of lentiviral plasmid, 15 μg of pSPAX2 (Addgene #8454) and 6 μg pMD2G (Addgene #12260) were mixed with 450 μL sterile water and 50 μL of 2.5M CaCl_2_ solution, previously filter sterilized. DNA mixture was added dropwise into 500 μL of 2X HeBS (50 mM HEPES, 10 mM KCl, 12 mM dextrose, 280 mM NaCl and 1.5 mM Na2PO4, pH 7.04) with constant vortexing. Solution was added dropwise on HEK 293T cells for 6h. Medium was replaced with 8mL of fresh 10% FBS DMEM medium for 48h. Supernatant was filtered through a 0.22 μm membrane and concentrated using a 100kDa cut-off Amicon centrigal unit. Aliquots were frozen at −80 °C.

For generation of CRISPR cells, single-guide RNA was selected using Benchling design tool, and are listed in Supplementary Table 3. Annealed oligonucleotides were cloned into pLentiCrispRv2-Puro as described by Sanjana and colleagues (https://doi.org/10.1038/nmeth.3047). Lentiviral particles were prepared as described above. hiPSCs were transduced in suspension and selected 48h post-transduction using 1 μg/mL of Puromycin.

### SDS-PAGE and Western blotting

Cells were washed with ice cold 1X PBS and lysed with 70 µL RIPA buffer (150 mM NaCl, 10 mM Tris–HCl pH 7.5, 1 mM EDTA, 1% Triton X-100, 0.1% SDS) supplemented with and 1X protease and phosphatase inhibitors. Mechanical disruption was performed by scrapping the cells off the plate, transferring the lysate into a 1.5mL Eppendorff tube and vortexing for 5 seconds followed by 5 min incubation on ice. Soluble proteins were collected by centrifugation at 16000 g (13200 rpm) for 10min at 4°C. Protein concentration was measured using Precision Red following the manufacturer’s instructions. Lysates in sample buffer were boiled for 5min.

For ppMLC blotting, cells were washed with 1X PBS and directly lysed using 250 μL of boiling 1X sample buffer. Lysates were sonicated 3 times for 10 seconds with 60 seconds incubation on ice between each blast. 30 μg of proteins were resolved on 1.5mm thick bis-tris acrylamide gels and transferred onto 0.2 μm nitrocellulose membrane using BioRad wet transfer system (50 V for 2hours). Membranes were blocked for > 30min at room temperature (RT) with 5% BSA in 1X TBS-T (10 mM Tris pH 8.0, 150 mM NaCl and 0.5% Tween 20) and incubated overnight at 4°C and gentle rocking with primary antibodies diluted in blocking buffer (Supplementary Table 2). Membranes were washed 3 times with 1X TBS-T and incubated for 1h at RT with AlexaFluor-conjugated secondary antibodies (Supplementary Table 2). After 3 additional washes in 1X TBS-T, membranes were scanned using Li-Cor Odyssey CLx system, analyzed and processed using Image StudioLite v. 5.2.

### Immunofluorescence

Cells were cultured on Matrigel-coated Ø 12mm coverglass, fixed with 4% paraformaldehyde for 10min, permeabilized for 5min at RT (20mM Glycine, 0.05% Triton X-100 in 1X PBS), and incubated with blocking buffer for > 30min (5% BSA in 1X PBS). Coverslips were transferred into a dark humidified chamber and incubated with primary and AlexaFluor-conjugated secondary antibodies (Supplementary Table 2) diluted in blocking buffer for 2 hours. Coverslips were washed in 1X PBS and mounted on glass slide using Fluoromount-G mounting solution. For ppMLC staining, samples were incubated overnight with primary antibody.

### Image acquisition, analysis and processing

Micrographs were obtained using an inverted Nikon A1-R confocal microscope equipped with a x20 objective (Numerical aperture NA 0.75), a x40 oil objective (NA 1.20) and a x100 oil objective (NA 1.40). Z-stacks covering the entire cell height were acquired. Maximum Intensity (MaxIP) Projections were obtained and images were analyzed and processed using Fiji (version 2.1.0/1.54f). Images processing did not include a denoising step except for Supplementary Video 3.

### MYPT1^CA^-NES-mNeonGreen cloning strategy and cell line generation

A gBlock was obtained from IDT, corresponding to truncated human MYPT1 (aa 1-300) in fusion with a PKI super Nuclear Export Signal and mNeonGreen sequence. The gBlock was amplified using Fusion polymerase and the following primers: Fwd 5’-CTGCTGACCGGTACCATGGCGGACGCGAAGC-3’ and Rev 5’-CATCATACGCGTCTACGATCCGCCACCGC-3’. PCR product and recipient backbone (pInducer10b-HA-KRas G12V Addgene #164928) were digested with AgeI-HF and MluI-HF for 1h at 37°C, gel purified, and ligated overnight at 16°C using T4 ligase. Ligation reactions were transformed into chemically competent E.Coli Stbl3 strain and selected on Ampicillin plates. Plasmids were isolated from single colonies (QIAprep Spin MiniPrep kit). The presence of the insert was confirmed by restriction digestion. Positive clones were sent for full plasmid sequencing provided by Genewiz Plasmid-EZ service before plasmid purification using Takara Bio NucleoBond XtraMidi kit. WT hiPSCs were transduced as described above and cells were selected 48h later with 1 μg/mL of Puromycin.

### Quantitative Polariztion Microscopy (QPOL)

The contractility of cells was measured using quantitative polarization microscopy (QPOL) as described previously ^33^. Briefly, QPOL was built on an inverted Axiovert microscope equipped with an Axiocam 506 color camera. A motorized linear polarizer (Thorlabs, Newton, NJ) and a circular polarizer were positioned in the illumination plane above the condenser and in the imaging plane, respectively. Images were captured using a 20x/0.5 N.A. polarization objective. For each field of view, image sequences were acquired with 10° intervals of the rotating polarizer over the range of 0–180° using Zen software. The polarized image sequences were then processed using a custom MATLAB code to generate pixel-by-pixel retardance maps. The obtained retardance images then underwent background subtraction and subsequent analysis with ImageJ. The retardance signal proportional to cell contractility was quantified as the average value of the background-subtracted retardance over the cell area.

### Flow cytometry analysis

T-mNeonGreen knock-in hESC were transduced with a lentivector expressing Histone 2B-mScarlet (pWPI-H2B-mScarlet). Cells were pre-treated overnight with Vehicle (water) or 10 μM Y-27632. Differentiation was initiated for 24 hrs as described above and cells were harvested as single cells. Briefly, cells were washed and gentle cell dissociation buffer was added for 8 min at 37°C. Cell suspension was centrifuged and resuspended in 500 μL PBS. Cells were passed through a Ø 40 μm strainer directly into 500 μL of 8% PFA (final concentration 4%) and fixed for 10 min on a wheel before sending them for cytometry using a 5-laser Fortessa analyzer (70 μm nozzle). Singlets were selected based of forward and side scatter. mScarlet (PE-Texas-RedA) and mNeonGreen (GFP-A) population were selected by drawing a gate on double negative WT hESC shifted by one order of magnitude. Percentage of mNeonGreen/mScarlet double positive was reported using the online flow cytometry analysis resource https://floreada.io.

### RNA isolation and RT-qPCR

Cells were washed and RNAs were isolated using RNeasy mini kit following manufacturer’s instructions. 1 μg of RNA was reverse transcribed to cDNA using the SuperScript III first-strand synthesis kit. cDNA was diluted 1:5 in water and mixed with Maxima SYBR Green/1mM of each primer (Supplementary Table 3). Quantitative PCR was performed on a BioRad CFX96 thermocycler with thermal cycling conditions as followed: Initial denaturation: 95°C for 10 min followed by 40 cycles of denaturation at 95°C for 15 sec, annealing at 58°C for 30 sec, extension at 72°C. Final step was performed for 10 sec at 95°C. C_t_ values from technical triplicates were averaged and normalized to GAPDH using the ΔΔC_t_ formula.

TCF expression levels (Supplementary Figure 7E) were curated from the transcriptomic data available on the Allen Institute website (https://www.allencell.org/genomics.html).

### Cell Fractionation

Following appropriate treatment, cells were washed with ice cold 1X PBS and collected as single cell suspension using gentle cell dissociation as previously described. Cell suspension was washed with cold 1X PBS and resuspended in 500 μL of cold PBS. 100 μL of the cell suspension was set aside and mixed with 30 μL of 4X sample buffer. Samples were sonicated for 15 sec at 20% power 3 times, boiled 5min and stored at −20 °C (Whole Cell Lysate fraction or WCL). Cell fractionation was performed using Cell signaling kit (#9038) and their protocol. Briefly, the remaining 400 μL of cell suspension was centrifuged at 500 x g at 4°C for 5min. Supernatant was removed and pellet was resuspended in 250 μL of CIB buffer, vortexed for 5 sec and incubated on ice for 5 min. Samples were centrifuged at 500 x g at 4°C for 5min and supernatant was stored in pre-chilled tube (= Cytoplasmic fraction). Pellet was resuspended in 250 μL of MIB buffer, vortexed for 15 sec and incubated on ice for 5 min. Samples were centrifuged at 8000 x g at 4°C for 5min and supernatant was discarded (= Membrane and organelle fraction). Pellet was resupended in 125 μL of CyNIB buffer, sonicated for 5 sec at 20% power 3 times (cytoskeletal and nuclear fraction). Each fraction was mixed with the appropriate volume of 4X sample buffer to obtain a final concentration of 1X. Samples were boiled for 5min and centrifuged before loading 15 μL.

### Nuclear extraction and CUT & RUN Assay

#### Nuclei extraction

500,000 cells (per sample) were harvested using TrypLE, washed with 1X PBS, and centrifuged at 600 × g for 3 min at RT. The pellet was resuspended in Nuclear Extraction (NE) buffer (20 mM HEPES, pH 7.9, 10 mM KCl, 0.1% Triton X-100, 20% glycerol, 1 mM MnCl₂, 0.5 mM spermidine, 1X Roche Complete Protease Inhibitor EDTA-Free) and incubated for 10 min at 4 °C. After incubation, samples were centrifuged at 600 × g, and the isolated nuclei were resuspended in 100 μL of NE buffer per sample. Nuclei were incubated with 10 μL of magnetic Concanavalin A beads for 10 min at RT to facilitate binding.

#### DNA purification

Bead-bound nuclei were incubated with 3 μL of anti-β-catenin antibody in 50 μL of digitonin buffer (20 mM HEPES, pH 7.5, 150 mM NaCl, 0.5 mM spermidine, 0.01% digitonin, 1X Protease Inhibitor Cocktail) containing 2 mM EDTA overnight at 4 °C. Following incubation, samples were washed twice with digitonin buffer and resuspended in 50 μL of the same buffer supplemented with 1.5 μL of pAG-MNase. Samples were incubated for 10 min at RT, followed by two additional washes, and subjected to chromatin digestion at 4 °C for 2 hrs with 2 mM CaCl₂. To quench the reaction, 33 μL of STOP buffer (340 mM NaCl, 20 mM EDTA, 4 mM EGTA, 50 µg/mL RNase A, 50 µg/mL glycogen) was added, followed by incubation at 37 °C for 10 min. Beads were collected using a magnetic rack, and the supernatant containing released DNA was transferred to a new tube. DNA was purified using DNA purification buffers and spin columns following the manufacturer’s instructions and eluted in 30 μL of buffer

#### qPCR reaction & Data analysis

Reactions were prepared using 1 µL of purified DNA from the CUT and RUN, with 2 µL Forward and reverse primer mix (5 µM each), 5 µL SYBR Green master mix (Applied Biosystems), and 2 µL of nuclease-free water. Thermal cycling condition were set up on a BioRad CFX96 thermocycler as followed: Initial denaturation: 95°C for 10 min followed by 40 cycles of denaturation at 95°C for 15 sec, annealing at 60°C for 30 sec, extension at 72°C. Final step was performed for 10 sec at 95°C. Ct values from technical triplicates were averaged, and ΔCt was calculated by subtracting Ct IgG control from target. Fold enrichment was calculated using the ΔΔCt method: Fold enrichment=2^−ΔΔCt^.

### GFP-Trap immunoprecipitation

EGFP-CTNNB1 knock-in hiPSCs were harvested using TripLE and nuclei were isolated as described above. For each reaction, two wells were collected. Both 0h and 24h nuclei were cryopreserved by slowly freezing them in NE buffer using an isopropanol-filled chiller at −80°C. Nuclei were quickly thawed using a 37°C water bath, centrifuged at 600 x g and resuspended using 100 μL of lysis buffer (10 mM Tris/HCl pH 7.5, 150 mM NaCl, 0.5 mM EDTA, 0.5 % NP40). Nuclear lysates were incubated 5 min on ice and cleared by centrifugation (16000 x g for 10 min at 4°C. Soluble protein concentration was measured using Precision Red following the manufacturer’s instructions and pellets were mixed with 1X sample buffer (insoluble fraction). Meanwhile, GFP-Trap beads (20 μL/reaction) were equilibrated three times with 500 μL of ice-cold dilution buffer (10 mM Tris/HCl pH 7.5, 150 mM NaCl, 0.5 mM EDTA). 150-200 μg of soluble proteins were prepared in 500 μL of ice-cold dilution buffer and mixed with equilibrated GFP-Trap beads. Reactions were incubated for 2 hrs at 4°C on a wheel. Beads were then pelleted (2500 x g, 3min at 4°C) and washed three times with 500 μL of ice-cold wash buffer (10 mM Tris/HCl pH 7.5, 150 mM NaCl, 0.05 % NP40, 0.5 mM EDTA). Proteins were eluted by adding 50 μL of 2X sample buffer and boiled for 5 min. Eluted proteins were resolved by western blot as described above. TCF4 binding was measured by densitometry of the TCF4 band (prey) normalized to the GFP band (bait).

### Reproducibility and statistics

All datasets were analyzed using GraphPad Prism10 (version 10.0.3). Individual measurements (n) were averaged for each biological repeat (N). Datasets were tested for normality (Shapiro-Wilk test for normality of biological replicates) before applying the appropriate statistical test on the biological repeats (N), except if mentioned differently in the figure legend. Error bars represent S.D. except where stated otherwise. For datasets displayed as superplots, individual measurements and biological repeats are represented by small and large dots respectively. Violin plots display median (thick dotted line) and quartiles (thin dotted lines). Each dataset for a biological repeat is color coded. Significance levels are given as follows and exact p-values are indicated in each figure: n.s., not significant (*p* ≥ 0.05); \**p* < 0.05; \*\**p* < 0.01; \*\*\**p* < 0.001; and \*\*\*\**p* < 0.0001.

All experiments, including representative MaxIPs and western blots, were performed at least three times independently as biological repeats, unless stated otherwise in the legends. No statistical method was used to pre-determine the sample size, the experiments were not randomized and the investigators were not blinded to allocation during experiments and outcome assessment.

---

**Supplementary Figure 1:**
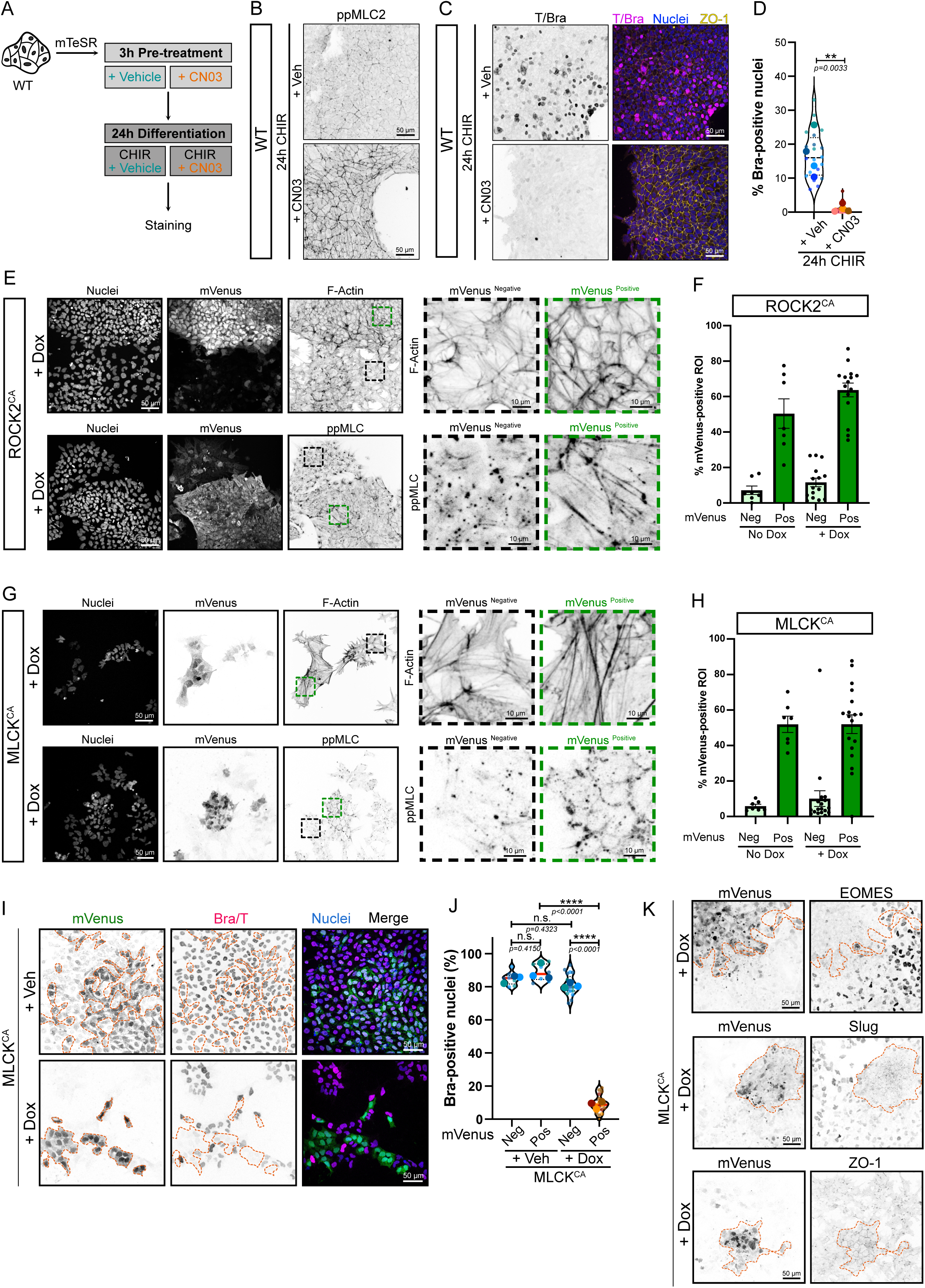
Increased actomyosin contractility is sufficient to block mesoderm commitment and EMT (Related to Figure 1) (A) Experimental design. hiPSCs were pre-treated with 4 µg/mL CN03 or vehicle for 3 hours. Differentiation was initiated with CHIR supplemented with Vehicle or 4 µg/mL CN03 for 24h before fixation and staining. (B) Representative MaxIP immunofluorescence of hiPSC treated as explained in (A) and stained for phospho-MLC2 (Inverted LUT). Scale bar = 50 µm (C, D) Representative MaxIP immunofluorescence of hiPSC treated as explained in (A) and stained for Brachyury (Inverted LUT), nuclei and ZO-1. Scale bar = 50 µm (C). Violin plot (median, quartiles) reporting the percentage of Brachyury-positive nuclei after 24h of differentiation and quantified across N=4 independent biological repeats (representing n=20 technical repeats). Unpaired t test was performed on biological replicates (D). (E) Representative MaxIP immunofluorescence of Doxycycline-induced ROCK2^CA^ hiPSCs (mVenus-positive) mixed with WT hiPSCs (mVenus-negative). Cells were stained for DNA (Nuclei), F-actin (top – Inverted LUT) and phospho T18/S19 Myosin Light Chain 2 (Bottom -ppMLC2 – Inverted LUT). Scale bar = 50 µm. Magnified views comparing mVenus-positive (Green dotted ROI) and mVenus-negative (Black dotted ROI) areas are shown for F-Actin and ppMLC2. Scale bar = 10 µm. (F) Percentage of the ROI area positive for mVenus based on cluster considered as mVenus-Negative and Positive using the ROCK2^CA^ line. On average, mVenus negative clusters have <15% of their area positive for mVenus. Mean and S.E.M are displayed. n=7 (no Dox) and n=15 fields of view (+Dox) across N=3 independent biological experiments. (G) Representative MaxIP immunofluorescence of Doxycycline-induced MLCK^CA^ hiPSCs (mVenus-positive) mixed with WT hiPSCs (mVenus-negative). Cells were stained for DNA (Nuclei), F-actin (top – Inverted LUT) and phospho T18/S19 Myosin Light Chain 2 (Bottom -ppMLC2 – Inverted LUT). Scale bar = 50 µm. Magnified views comparing mVenus-positive (Green dotted ROI) and mVenus-negative (Black dotted ROI) areas are shown for F-Actin and ppMLC2. Scale bar = 10 µm. (H) Percentage of the ROI area positive for mVenus based on cluster considered as mVenus-Negative and Positive using the MLCK^CA^ line. On average, mVenus negative clusters have <15% of their area positive for mVenus. Mean and S.E.M are displayed. n=7 (no Dox) and n=18 fields of view (+Dox) across N=3 independent biological experiments. (I, J) Representative MaxIP immunofluorescence for Brachyury in Vehicle (+Veh) and Doxycycline-induced MLCK^CA^ co-culture. mVenus-positive cell clusters are highlighted by an orange dotted line. Individual channels are inverted LUT. Scale bar = 50 µm (I). Percentage of Brachyury-positive nuclei is reported as violin plot, comparing mVenus-positive *vs* mVenus-negative cluster in the presence or absence of Doxycycline. Median (Plain red line) and quartiles (Dotted black lines). n=7 (+Veh) and n=18 (+ Dox) fields of view (small dots) across N=3 independent biological repeats (large dots). One-way ANOVA with Tukey’s multiple comparisons post-test was performed (J). (K) Representative MaxIP immunofluorescence for EOMES, Slug and ZO-1 in Doxycycline-induced MLCK^CA^ co-culture. mVenus-positive cell clusters are highlighted by an orange dotted line. Individual channels are inverted LUT. Scale bar = 50 µm.

**Supplementary Figure 2:**
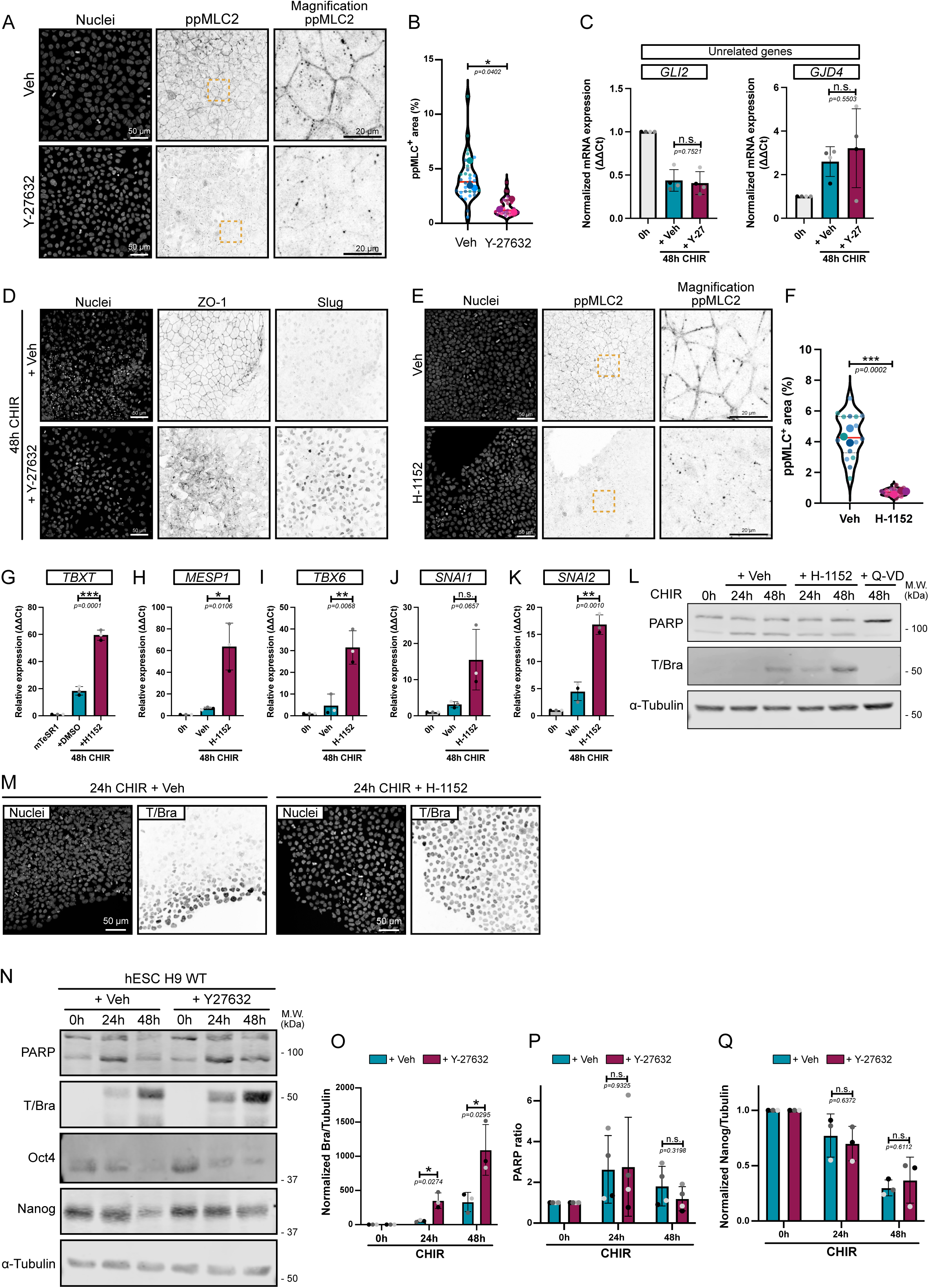
Pharmaceutical inhibition of actomyosin contractility promotes hiPSC and hESC conversion to the mesoderm lineage (Related to Figure 3) (A, B) Representative MaxIP immunofluorescences of hiPSCs treated with a ROCK inhibitor (Y-27632) or Vehicle (H_2_O) and stained for nuclei marker (DNA) and phospho T18/S19 Myosin Light Chain 2 (ppMLC2 – Inverted LUT). Scale bar = 50 µm. Magnified views of the yellow dotted ROI are shown for ppMLC2. Scale bar = 20 µm (A). Fraction of the cellular area positive for ppMLC2 following Y-27632 treatment and reported as violin pots. Median (Plain red line) and quartiles (Dotted black lines). n=30 technical repeats across N=3 independent biological repeats. Two-tailed unpaired t test was performed on the biological repeat (B). (C) Relative expression of genes not directly related to mesoderm commitment at 0h and 48h post CHIR treatment in the presence (+Y-27) or absence (+Veh) of ROCK inhibitor. N=4 independent biological repeats. Mean and S.D. are displayed. Two-tailed unpaired t test was performed. (D) Representative MaxIP immunofluorescences of hiPSC treated 48h with CHIR in the presence (+ Y-27632) or absence (+ Veh) of ROCK inhibitor. Cells were stained for nuclei marker (DNA), EMT markers (ZO-1 and Slug – Inverted LUT). Scale bar = 50 µm. (E, F) Representative MaxIP immunofluorescences of hiPSCs treated with a ROCK inhibitor (H-1152) or Vehicle (H_2_O) and stained for nuclei marker (DNA) and phospho T18/S19 Myosin Light Chain 2 (ppMLC2 – Inverted LUT). Scale bar = 50 µm. Magnified views of the yellow dotted ROI are shown for ppMLC2. Scale bar = 20 µm (E). Fraction of the cellular area positive for ppMLC2 following H-1152 treatment and reported as violin pots. Median (Plain red line) and quartiles (Dotted black lines). n=15 technical repeats across N=3 independent biological repeats. Two-tailed unpaired t test was performed on the biological repeats (F). (G-K) Relative expression of mesoderm markers (*TBXT*, *MESP1, TBX6*) (G-I) and EMT markers (*SNAI1, SNAI2*) (J, K) 48h post CHIR treatment in the presence (+H-1152) or absence (+Veh) of ROCK inhibitor. N=3 independent biological repeats. Mean and S.D. are displayed. Two-tailed unpaired t test was performed. (L) Immunoblot probing for mesoderm marker expression (T/Bra) and cell death marker (PARP cleavage) during hiPSC-to-mesoderm commitment (0h-48h CHIR) in the presence of ROCK inhibitor (+H-1152). Addition of the pan-caspase inhibitor Q-VD-OPH (+ Q-VD) to the differentiation medium totally blocks mesoderm commitment and was used as control. α-Tubulin was used as loading control. Molecular weights (M.W.) are displayed on the right side. (M) Representative MaxIP immunofluorescences for mesoderm marker Brachyury (T/Bra – Inverted LUT) and nuclei. Cells were fixed 24h post-CHIR treatment in the presence or absence of ROCK inhibitor H-1152. Scale bar = 50 µm (N) Representative immunoblot of PARP (cell death), T/Bra (mesoderm), Oct4 and Nanog (Pluripotency markers) using hESC H9 in the presence (+ Y-27632) or absence (+ Veh) of ROCK inhibitor, as shown in Figure 3A. Molecular weights (M.W.) are displayed on the right side. (O-Q) Brachyury expression (O), PARP cleavage (P) and Nanog expression (Q) was quantified by densitometry and normalized to α-Tubulin as loading control across N=4 (PARP) and N=3 (Brachyury and Nanog) independent biological repeats. Mean and S.D. are displayed. Two-tailed unpaired t test was performed.

**Supplementary Figure 3:**
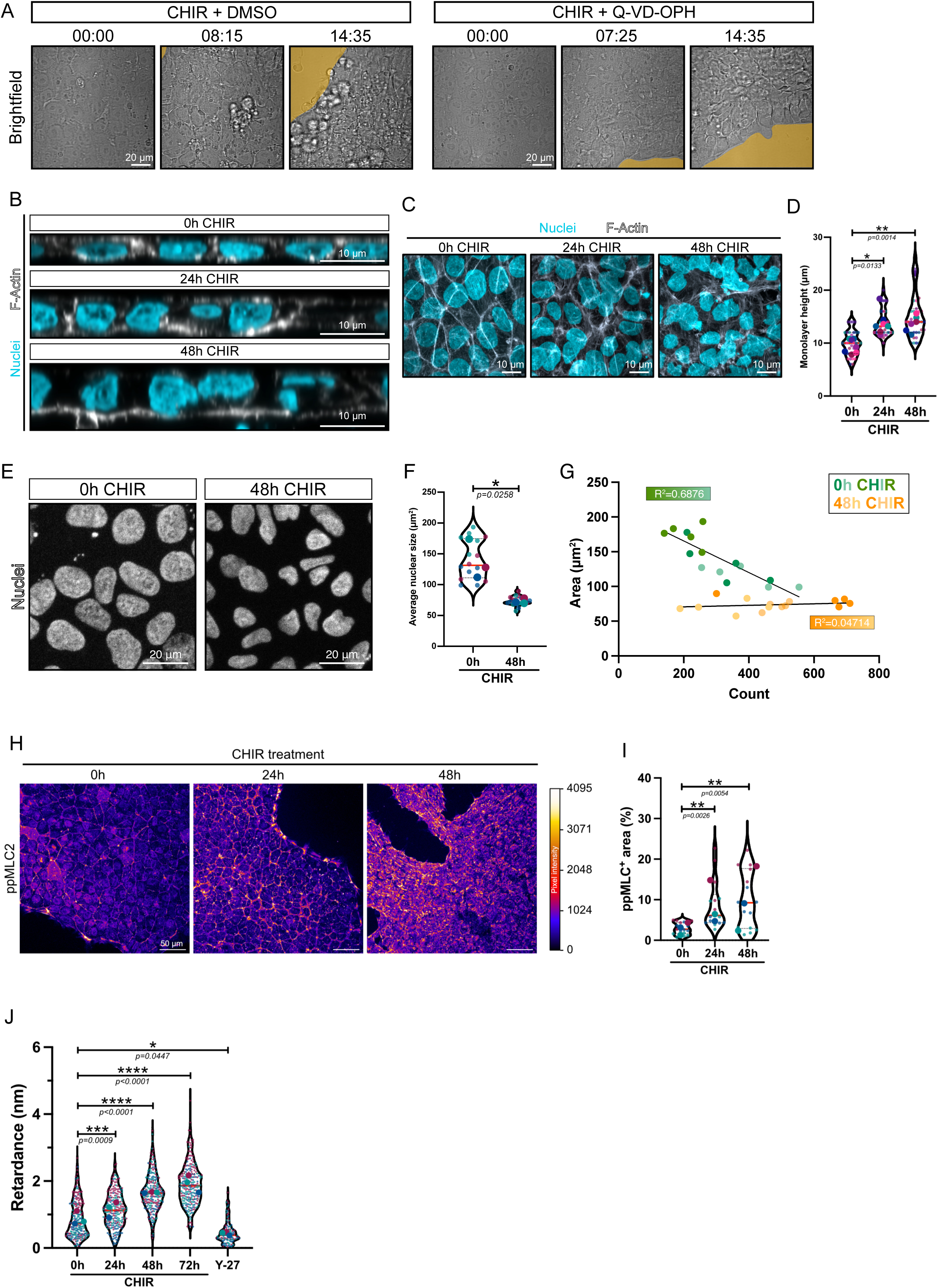
hiPSC differentiation along the cardiac mesoderm increases actomyosin-driven contractility. (A) Bright field imaging of WT hiPSCs treated with CHIR in the presence or absence of a pan-Caspase inhibitor Q-VD-OPH. Retraction area is highlighted in yellow. Time is presented as hh:mm. Scale bar = 20 µm. (Related to Supplementary movie 1). (B, C) Confocal XZ orthogonal views (B) and Maximum Intensity Projections (MaxIP) (C) of hiPSCs at basal state (0h CHIR) or post-CHIR treatment (24h and 48h CHIR), stained for DNA (Cyan) and F-Actin (Grey). Scale bar = 10 µm. (D) Quantification of the monolayer height at 0h, 24h and 48h post CHIR treatment from (B). Violin plots represent individual measurements (small dots) averaged for each biological repeat (large dots). Median (Plain red line) and quartiles (Dotted black lines). n=32 technical repeats across N=7 independent biological repeats. A Kruskal-Wallis test with Dunn’s multiple comparison post-test was performed on biological repeats. (E-G) Representative MaxIP immunofluorescence of WT hiPSC during CHIR treatment. Fixed cells were stained for DNA marker (Hoechst). Scale bar = 20 µm (E). Average nuclear size from n=15 fields of view across N=3 independent biological repeats. Median (Plain red line) and quartiles (Dotted black lines) are displayed. Two-tailed unpaired t test was performed (F). Correlation of nuclear area over cell number for each field of view during CHIR treatment. Biological repeats are color-coded. Pearson correlation coefficient is displayed for each treatment (G). (H) Representative MaxIP immunofluorescence of hiPSCs at basal state (0h) or post-CHIR treatment (24 and 48h) stained for phospho T18/S19 MLC2 (ppMLC2). Pixels are color-coded by intensity using the fire LUT. Scale bar = 50 μm (I) Percentage of ppMLC2-positive area at 0hr, 24hrs, 48hrs post CHIR is reported as violin plots. Median (Plain red line) and quartiles (Dotted black lines). n=15 technical repeats across N=3 independent biological repeats. A Kruskal-Wallis test with Dunn’s multiple comparison post-test was performed on the technical repeats. (J) Retardance measurements obtained from hiPSCs at basal state (0h), post-CHIR treatment (24h, 48h, 72h) or treated with Y-27632 at 0h (Y-27) using Quantitative Polarization Microscopy imaging. Median (Plain red line) and quartiles (Dotted black lines). n=240 technical repeats for 0hr, 24hrs, 48hrs, 72hrs and n=120 technical repeats for Y-27 across N=3 independent biological repeats. A one-way ANOVA with Šidák’s multiple comparisons post-test was performed on the biological repeats.

**Supplementary Figure 4:**
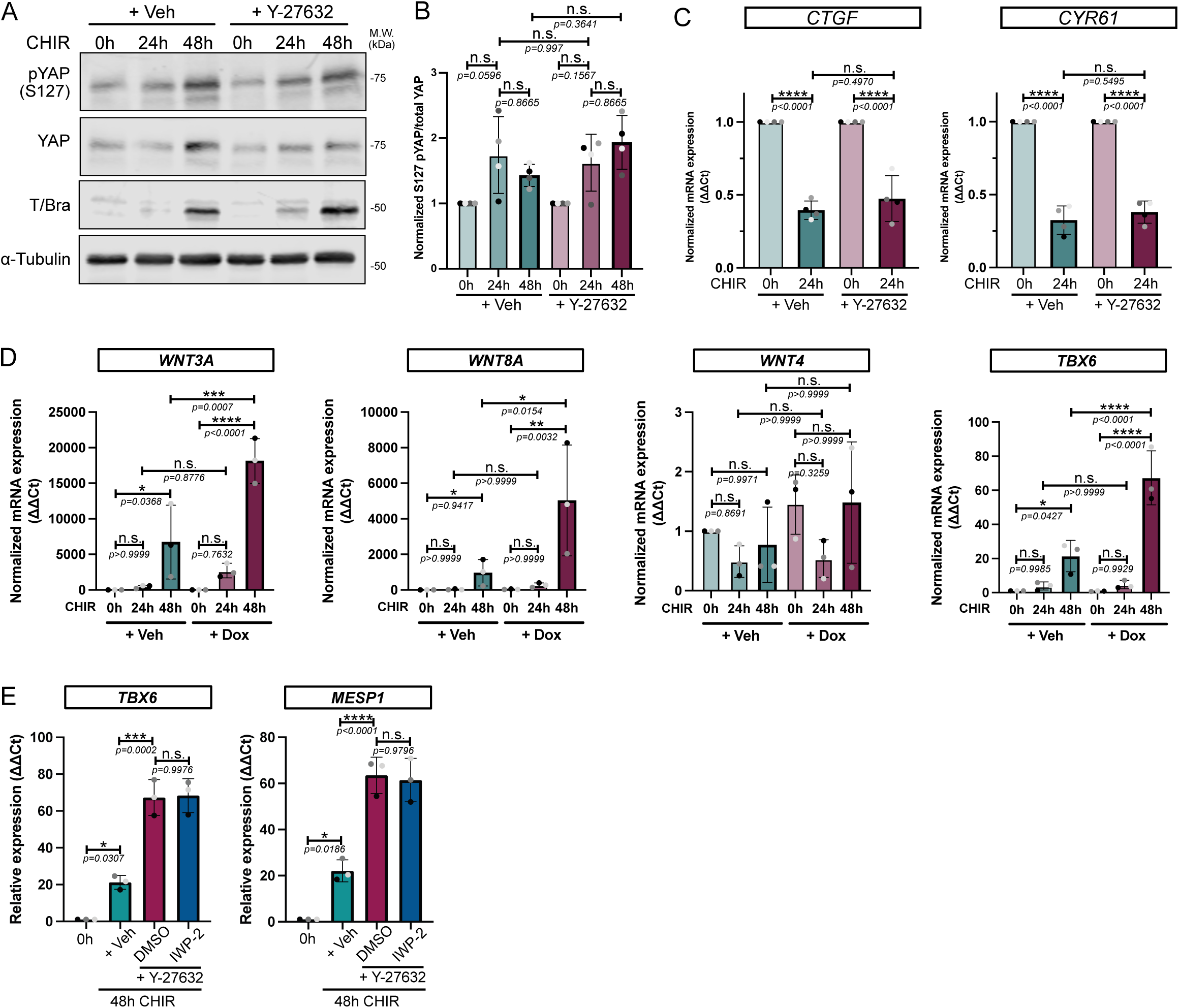
Key mechanosensitive pathways are not involved in mesoderm specification enhancement following decreased contractility. (A, B) Immunoblot for phospho S127 YAP (pYAP) and total YAP following differentiation (0-48h CHIR) in the presence or absence of ROCK inhibitor Y-27632. Brachyury expression was also probed as an internal positive control and α-Tubulin was used as loading control. Molecular weights (M.W.) are displayed on the right side (A). Ratio of phospho/total YAP was quantified across N=4 independent biological repeats. Mean and S.D. are displayed. One-way ANOVA with Šidák’s multiple comparisons post-test was performed (B) (C) Relative expression of YAP-target genes *CTGF* and *CYR61* during differentiation +/-Y-27632. N=4 independent biological repeats. Mean and S.D. are displayed. One-way ANOVA with Šidák’s multiple comparisons post-test was performed. (D) Relative expression of canonical WNT (*WNT3A, WNT8A*), non-canonical WNT (*WNT4*) and mesoderm marker (*TBX6*) in MYPT1^CA^-NES-mNG cells treated with CHIR (0-48h) in the presence of Vehicle or Doxycycline. N=3 independent biological repeats. Mean and S.D. are displayed. One-way ANOVA with Šidák’s multiple comparisons post-test was performed. (E) Relative expression of *TBX6* and *MESP1* (mesoderm markers) following CHIR treatment +/-Y-27632 complemented or not with 7.5 µM Porcupine inhibitor (IWP-2 or DMSO respectively). N=3 independent biological repeats. Mean and S.D. are displayed. One-way ANOVA with Tukey’s multiple comparisons post-test was performed.

**Supplementary Figure 5:**
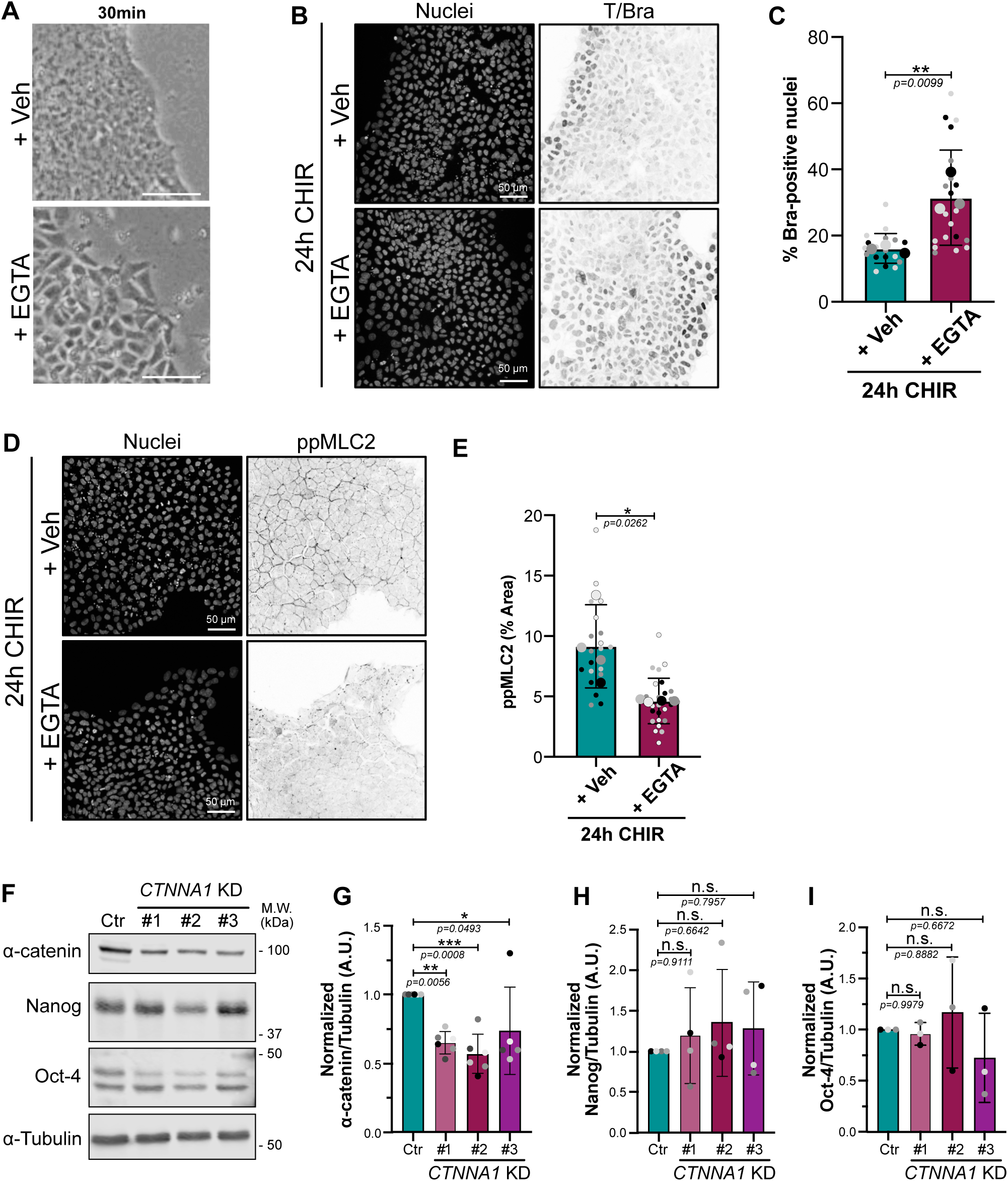
EGTA treatment and validation *CTNNA1* knockdown hiPSCs (related to Figure 4) (A) Bright field picture showing the colony morphology following EGTA pre-treatment. Colony appears less compact and individual cell boundaries can be distinguished. Scale bar = 20 μm. (B, C) Representative immunofluorescence of hiPSCs treated as shown in Figure 4A and stained for Brachyury. Scale bar = 50 μm (B). Quantification of Brachyury-positive nuclei is shown following EGTA treatment. Mean and S.D. are displayed. n=18 (Veh) and n=20 fields of view across N=3 independent biological repeats. Two-tailed unpaired t test was performed on biological repeats (C). (D, E) Representative immunofluorescence of hiPSC treated as shown in Figure 4A and stained for ppMLC2. Scale bar = 50 μm (D). Quantification of ppMLC2-positive area is shown following EGTA treatment. Mean and S.D. are displayed. n=20 (Veh) and n=25 fields of view across N=4 independent biological repeats. Two-tailed unpaired t test was performed on biological repeats (E). (F-I) Representative immunoblot of control (Ctr) or *CTNNA1* knockdown hiPSCs, probed for α-catenin, Nanog and Oct-4 (pluripotency) and α-Tubulin as loading control. Molecular weights (M.W.) are displayed on the right side (F). Quantification of α-catenin (G), Nanog (H) and Oct-4 (I) expression was obtained by densitometry across N=5-6 (α-catenin), N=4 (Nanog) and N=3 (Oct-4) independent biological repeats. Mean and S.D. are displayed. One-Way ANOVA with Dunnett’s multiple comparisons post-test was performed.

**Supplementary Figure 6:**
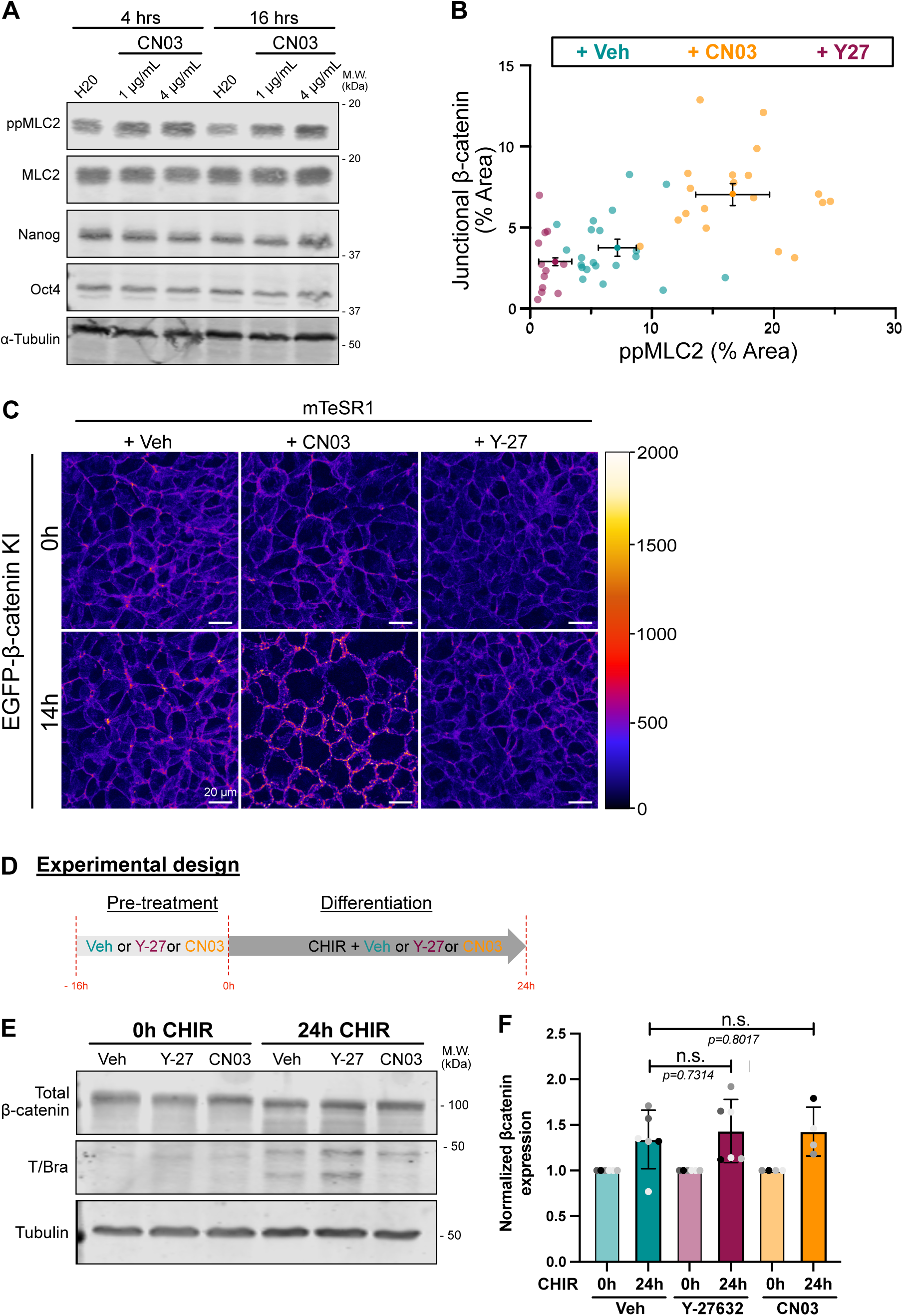
Contractility affects β-catenin localization at adherens junctions (related to Figure 5) (A) Immunoblot assessing the effects of CN03 treatment at 4h and 16h. Membrane was probed for total and phospho-ML2 (MLC2 and ppMLC2 respectively), Nanog and Oct-4 and α-Tubulin as loading control. Molecular weights (M.W.) are displayed on the right side. (B) Independent technical repeats from Figure 5 A-B. Repeats across the 3 biological repeats were averaged and shown as a dot with error bars (S.D.). (C) Still picture from overnight confocal imaging of mEGFP-β-catenin knock-in hiPSC treated with Vehicle, CN03 and Y-27632 at basal state. Pixels are color-coded by intensity using the Red Fire LUT. Scale bar = 20 µm. (Related to Supplementary video 2). (D) Experimental design. Cells were pre-treated overnight with Vehicle or Y-27632 or CN03 in mTeSR1 and differentiated using CHIR media complemented with Vehicle or Y-27632 or CN03 for 24h before collecting protein lysates (E, F) Representative immunoblot for total β-catenin. Membrane was also probed for Brachyury to show effect of each drug and α-Tubulin as loading control. Molecular weights (M.W.) are displayed on the right side (E). β-catenin expression was quantified across N=6 (Vehicle and Y-27632) and N=4 (CN03) independent biological repeats. Mean and S.D. are displayed. One-way ANOVA with Šidák’s multiple comparisons post-test was performed (F).

**Supplementary Figure 7:**
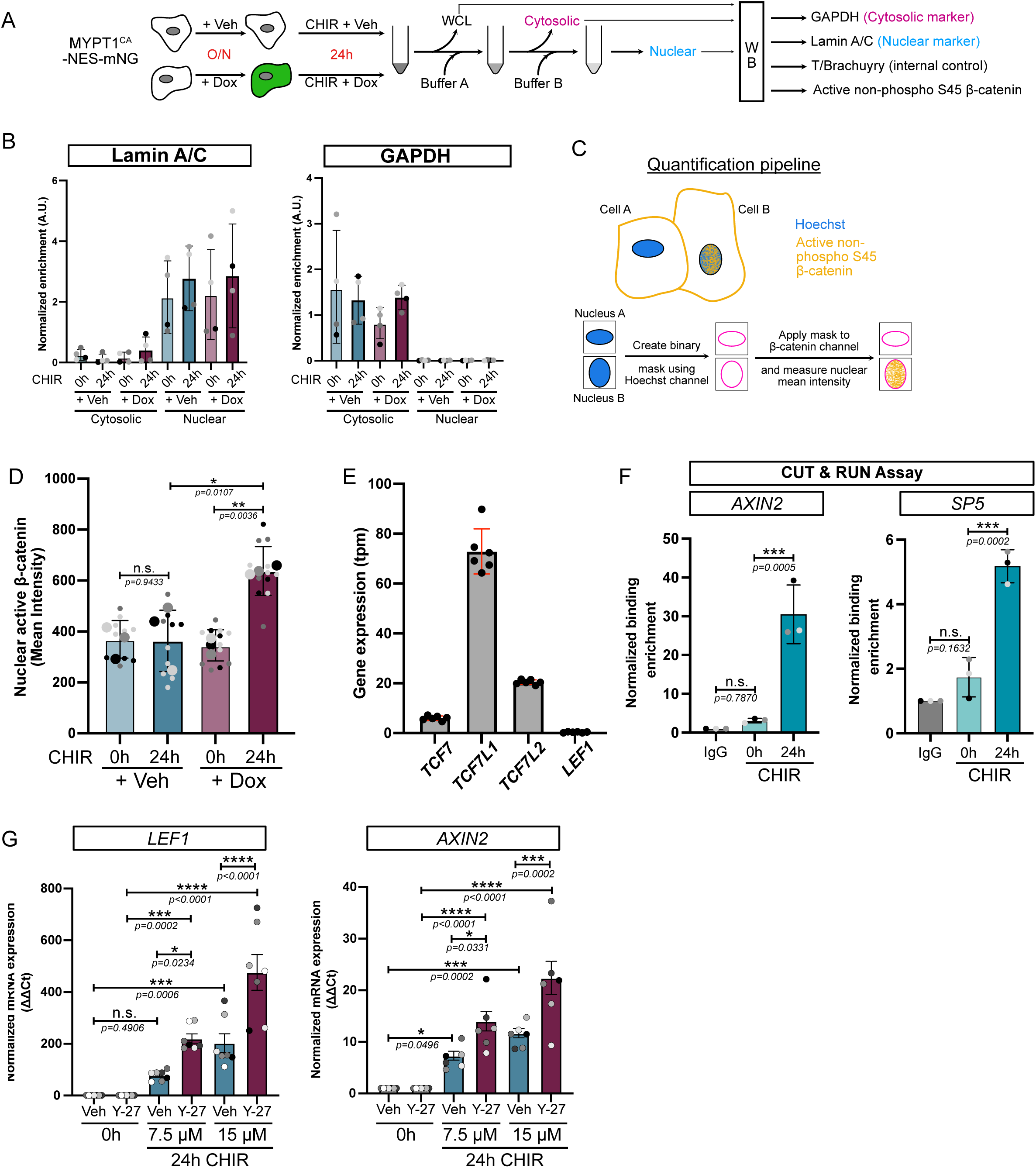
Reduced cell contractility promotes nuclear accumulation of β-catenin (related to Figure 6) (A) Experiment design for treatments and cell fractionation. (B) Compartment purity following fractionation was assessed by quantifying Lamin A/C and GAPDH enrichment in both, cytosolic and nuclear fractions across N=4 independent biological repeats. (C) Schematic representation of the quantification pipeline for nuclear β-catenin. (D) Quantification of active non-phospho S45 β-catenin mean intensity as depicted in (C). n = 10-12 technical repeats across N=3 independent biological repeats. Mean and S.D. are displayed. One-way ANOVA with Šidák’s multiple comparisons post-test was performed. (E) Expression of TCF genes (transcripts per million – tpm) from parental WTC hiPSC obtained from the Allen Institute transcriptomic data. N=6 independent biological repeats. (F) Positive control for CUT & RUN assay using IgG and β-catenin binding to WNT target genes (*AXIN2* and *SP5*) between 0h and 24h of CHIR treatment. Mean and S.D. are displayed. N=3 independent biological repeats. One-Way ANOVA with Dunnett’s multiple comparisons post-test was performed. (G) Relative expression of WNT target genes (*LEF1* and *AXIN2*) with increasing concentration of CHIR in the presence (Magenta) or absence (Cyan) of ROCK inhibitor Y-27632. N=7 independent biological repeats. Mean and S.E.M. are displayed. One-way ANOVA with Šidák’s multiple comparisons post-test was performed.

<u>Supplementary Video 1</u>

Bright field movie of hiPSC colony treated with CHIR supplemented with Vehicle (DMSO) or Q-VD-OPH (cell death inhibitor). Time in hh:mm. Scale bar = 20 μm.

<u>Supplementary Video 2</u>

Confocal imaging of mEGFP-β-catenin knock-in hiPSC treated with Vehicle (water), 4μg/mL CN03 or 10 μM Y-27632 in mTeSR1. Red Fire LUT reflects the pixel intensity. Time in hh:mm:ss. Scale bar = 20 μm.

<u>Supplementary Video 3</u>

Confocal imaging of mEGFP-β-catenin knock-in hiPSC treated with CHIR supplemented with Vehicle (water) or 10 μM Y-27632. Time in hh:mm. Scale bar = 10 μm.

